# Longitudinal analysis of diffuse glioma reveals cell state dynamics at recurrence associated with changes in genetics and the microenvironment

**DOI:** 10.1101/2021.05.03.442486

**Authors:** Frederick S Varn, Kevin C Johnson, Taylor E Wade, Tathiane M Malta, Thais S Sabedot, Floris P Barthel, Hoon Kim, Nazia Ahmed, Indrani Datta, Jill S Barnholtz-Sloan, Spyridon Bakas, Fulvio D’Angelo, Hui K Gan, Luciano Garofano, Jason T Huse, Mustafa Khasraw, Emre Kocakavuk, Simona Migliozzi, D. Ryan Ormond, Sun Ha Paek, Erwin G Van Meir, Annemiek M.E. Walenkamp, Colin Watts, Michael Weller, Tobias Weiss, Pieter Wesseling, Lucy F Stead, Laila M Poisson, Houtan Noushmehr, Antonio Iavarone, Roel GW Verhaak, The GLASS Consortium

## Abstract

To interrogate the factors driving therapy resistance in diffuse glioma, we collected and analyzed RNA and/or DNA sequencing data from temporally separated tumor pairs of 292 adult patients with IDH-wild-type or IDH-mutant glioma. Tumors recurred in distinct manners that were dependent on IDH mutation status and attributable to changes in histological feature composition, somatic alterations, and microenvironment interactions. Hypermutation and acquired *CDKN2A* deletions associated with an increase in proliferating stem-like malignant cells at recurrence in both glioma subtypes, reflecting active tumor growth. IDH-wild-type tumors were more invasive at recurrence, and their malignant cells exhibited increased expression of neuronal signaling programs that reflected a possible role for neuronal interactions in promoting glioma progression. Mesenchymal transition was associated with the presence of a specific myeloid cell state defined by unique ligand-receptor interactions with malignant cells. Collectively, our results uncover recurrence-associated changes that could be targetable to shape disease progression following initial diagnosis.

## Introduction

Diffuse gliomas in adults are aggressive primary tumors of the central nervous system that are characterized by a poor prognosis and the development of resistance to a treatment regimen that typically includes surgery, alkylating chemotherapy, and radiotherapy (Stupp et al., 2005; Wen et al., 2020). Genomic profiling of diffuse glioma has identified genomic drivers of disease progression and led to the definition of clinically relevant subtypes based on the presence of somatic mutations in the isocitrate dehydrogenase (IDH) genes and co-deletion of chromosome arms 1p and 19q (Cancer Genome Atlas Research et al., 2015; Ceccarelli et al., 2016; Eckel-Passow et al., 2015; Louis et al., 2016; Weller et al., 2015; Yan et al., 2009). Transcriptional profiling of whole tumors and single cells has revealed that the gene expression programs in malignant glioma cells are influenced by underlying somatic alterations and interactions with the tumor microenvironment. Additionally, malignant cells exhibit high plasticity that enables them to respond dynamically to diverse challenges (Johnson et al., 2020; Neftel et al., 2019; Patel et al., 2014; Phillips et al., 2006; Venteicher et al., 2017; Verhaak et al., 2010; Wang et al., 2017). Studies of changes relating to therapy using bulk genomics approaches have revealed mesenchymal transitions and both branching and linear evolutionary patterns (Barthel et al., 2019; Kim et al., 2015a; Kim et al., 2015b; Korber et al., 2019; Wang et al., 2016; Wang et al., 2017). However, the extent to which individual malignant glioma and immune cells interact and evolve over time to facilitate therapy resistance remains poorly understood.

To identify the drivers of treatment resistance in glioma, we established the Glioma Longitudinal Analysis Consortium (GLASS) (Bakas et al., 2020; Barthel et al., 2019; Consortium, 2018). In our initial effort, we assembled a set of longitudinal whole-exome and whole-genome sequencing data from 222 patients to define the clonal dynamics that allow each glioma subtype to escape therapy. In the current study, we build upon these analyses by integrating this genomic dataset with overlapping and complementary longitudinal transcriptomic data. We apply single-cell-based deconvolution approaches to these data to infer a tumor’s physical structure and identify the cell state interactions across IDH-wild-type and IDH-mutant glioma. Collectively, we find that gliomas exhibit several common transcriptional and compositional changes at recurrence that represent promising therapeutic targets for delaying disease progression.

## Results

### Overview of the GLASS Cohort

We expanded the GLASS cohort with an emphasis on collecting orthogonal RNA sequencing profiles to include data from a total of 351 patients treated across 35 hospitals (**Table S1**). After applying genomic and clinical quality control filters, the resulting dataset included genomic data from a total of 292 patients, with 150 having RNA sequencing data available for at least two time points, 243 having DNA sequencing data available for at least two time points, and 101 having overlapping RNA and DNA available at each time point. The cohort of 150 tumors used for RNA sequencing analyses comprised each of the three major glioma subtypes, with 114 IDH wild-type (IDH-wild-type), 27 IDH mutant 1p/19q intact (IDH-mutant-noncodel), and 9 IDH mutant 1p/19q co-deleted (IDH-mutant-codel) glioma pairs (**Figure 1A**). Given the limited number of IDH-mutant- codel cases, we grouped the IDH-mutant categories, unless specified otherwise. To facilitate further investigation and discovery of the drivers of treatment resistance in glioma, we have made this resource available to the research community (https://www.synapse.org/#!Synapse:syn21589818).

**Figure 1.**
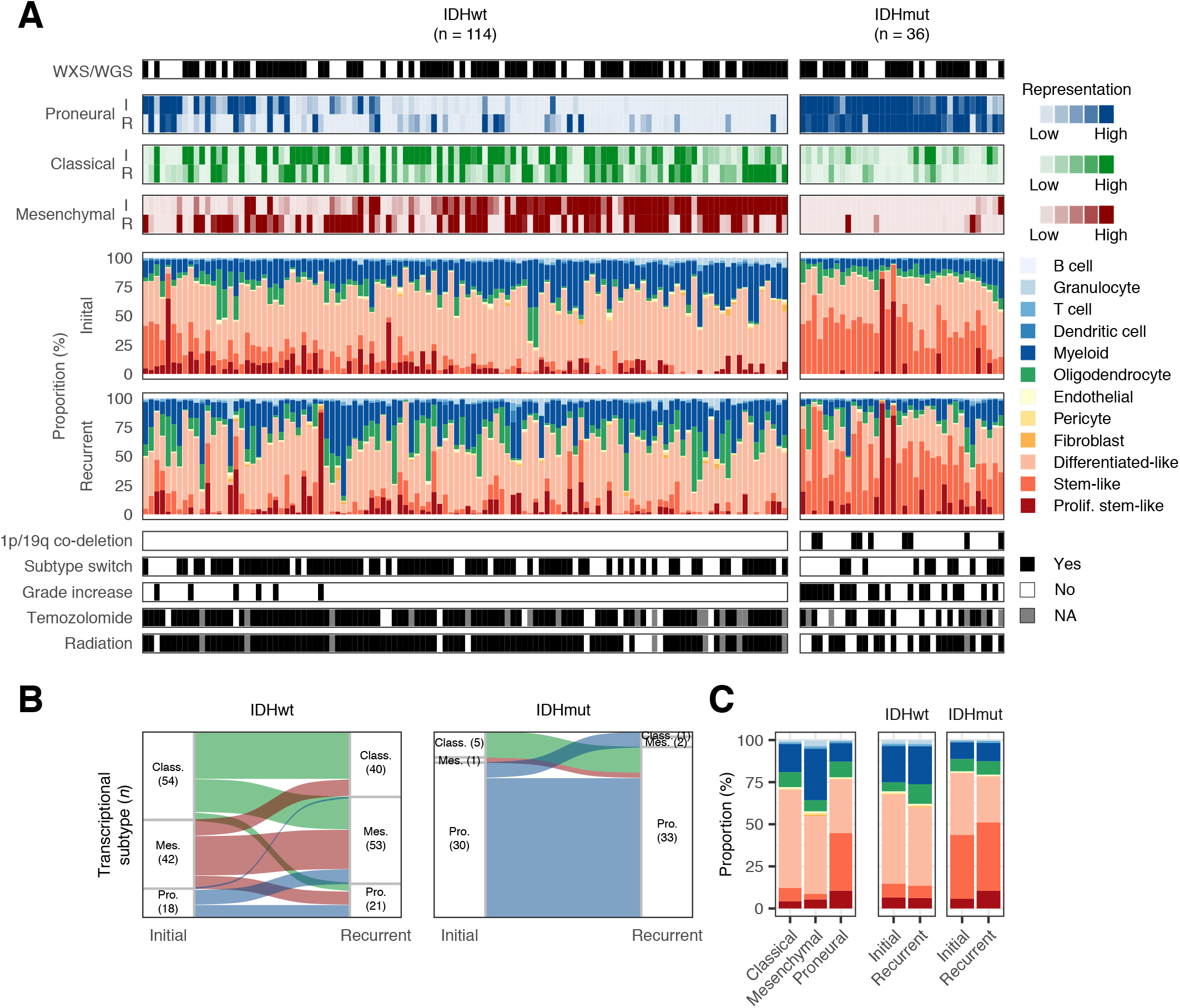
Diffuse glioma exhibits transcriptional and cellular heterogeneity across samples, subtypes, and time. (A) Overview of the GLASS dataset. Each column represents a tumor pair, and their initial (I) and recurrent (R) samples are labelled. All tumor pairs with RNAseq data at each time point are included. Pairs are arranged based on the representation of the proneural and mesenchymal subtypes in their initial tumors. The first track indicates whether there is whole exome or whole genome sequencing data available for that pair. The next three tracks indicate the representation of each bulk subtype across each sample. The stacked bar plots indicate the cell state composition of each sample based on the single cell-based deconvolution method, CIBERSORTx. The bottom tracks indicate molecular and clinical information for each tumor pair. (B) Sankey plot indicating whether the highest-scoring transcriptional subtype changed at recurrence. Each color reflects the transcriptional subtype in the initial tumors. Number in parentheses indicates number of samples of that subtype. (C) Left: The average cell state composition of each bulk transcriptional subtype for all initial GLASS tumors. Right: The average cell state composition of initial and recurrent tumors stratified by IDH mutation status. Colors in (C) are identical to those used in (A).

### Transcriptional activity and cellular composition in glioma is variable over time

To obtain a baseline understanding of transcriptional evolution in glioma, we assessed the representation of the classical, mesenchymal, and proneural transcriptional subtypes in each sample. IDH-wild-type tumors exhibited primarily classical and mesenchymal characteristics compared to IDH-mutant tumors, which were largely proneural (**Figure 1A**). Longitudinally, the dominant subtype in IDH-wild-type tumors switched in 46% of patients, with classical to mesenchymal being the most common transition. IDH-mutant tumors were more stable, with 75% of tumors remaining proneural at both time points (**Figure 1B**). Classical IDH-wild-type and IDH-mutant tumors switched subtype 50% of the time, resulting in an overall reduction of classical tumors at recurrence. The occurrence of this transition was significant (*P* = 0.04, Fisher’s exact test), suggesting that the tumor cells underlying the classical subtype may have higher plasticity than other subtypes.

To understand the cellular phenotypes underlying the transcriptional dynamics over time, we deconvoluted the GLASS gene expression dataset using CIBERSORTx (Newman et al., 2019) integrated with reference cell state signatures derived from our previously established collection of 55,284 single-cell transcriptomes from 11 adult patients spanning glioma subtypes and time points (Johnson et al., 2020) (**Table S2, Table S3**). Unsupervised analyses of the single-cell data had previously identified 12 cell states that represented the glial, stromal, immune, and malignant compartments commonly present in glioma. The malignant population expressed a shared set of markers (e.g., *SOX2*) and was split across three pan-glioma cell states, differentiated-like, stem-like, and proliferating stem-like, that together capture the gradient between development, lineage commitment, and proliferative status that has been observed across numerous glioma single-cell studies (Bhaduri et al., 2020; Castellan et al., 2021; Couturier et al., 2020; Garofano et al., 2021; Neftel et al., 2019; Richards et al., 2021; Tirosh et al., 2016; Venteicher et al., 2017; Wang et al., 2019; Yuan et al., 2018). Specifically, the differentiated-like state encompassed malignant cells exhibiting oligodendrocyte-like, astrocyte-like, and mesenchymal-like processes, while the stem-like states could be segregated by cell cycle activity and resembled undifferentiated and progenitor-like malignant cells (Neftel et al., 2019; Venteicher et al., 2017). To validate this approach, we applied CIBERSORTx to 1) a series of synthetic mixtures composed of single cells from our reference dataset that had been left out of the signature creation process; and 2) bulk RNAseq profiles from our reference dataset that had their true proportions determined from scRNAseq (**Figure S1A** and **S1B**).

When applying our deconvolution approach to the GLASS dataset, we observed variations in cellular composition across each subtype consistent with prior literature (Neftel et al., 2019; Wang et al., 2017). Classical and mesenchymal tumors had high levels of differentiated-like malignant cells, with the latter also having high levels of stromal and immune cells, and proneural tumors had high levels of proliferating stem-like and stem-like malignant cells (**Figure 1C**). Longitudinally, we found that IDH-wild-type tumors had significantly higher levels of oligodendrocytes and significantly lower levels of differentiated-like malignant cells at recurrence (*P* = 2e-5 and 2e-3, paired t-test). These changes remained significant even when accounting for differences in the surgical resection extent at each time point, suggesting a greater admixture of malignant cells and oligodendrocytes (**Figure S1C**). We observed similar changes in cellular composition when using an independently published integrative model of cell state classification that has been established for IDH-wild-type glioma, including a significant decrease at recurrence in the astrocyte-like malignant cell state that is dominant in classical IDH-wild-type tumors (*P* = 2e-3, paired t-test; **Figure S1D**) (Neftel et al., 2019). Recurrent IDH-mutant tumors exhibited significantly higher levels of proliferating stem-like malignant cells and significantly lower levels of differentiated-like malignant cells (*P* = 3e-3 and 2e-5, paired t-test; **Figure 1C**). Stratifying this group by 1p/19q co-deletion status revealed that the increase in proliferating stem-like cells was only significant in IDH-mutant-noncodels, while IDH-mutant-codels exhibited a significant increase in stem-like cells (*P* = 0.04, paired t-test; **Figure S1E**). Overall, the differences IDH-wild-type and IDH-mutant tumors exhibited over time suggested that distinct factors influence recurrence in each subtype.

### Histological features underlie subtype switching and cell state changes at recurrence

Intratumoral heterogeneity is a hallmark of glioma and is abundant in hematoxylin and eosin-stained tissue slides, where features such as microvascular proliferation and necrosis are used for diagnosis and grading by pathologists (Hambardzumyan and Bergers, 2015; Kristensen et al., 2019). The Ivy Glioblastoma Atlas Project has defined and microdissected five “anatomic” features on the basis of reference histology: 1) the leading edge of the tumor, 2) the infiltrating tumor front, 3) the cellular tumor, 4) pseudopalisading cells around necrosis, and 5) microvascular proliferation (Puchalski et al., 2018). They have shown that each of these features has a distinct transcriptional profile, suggesting that changes in a tumor’s cell state composition at recurrence reflect changes in a tumor’s underlying physical structure. To obtain a better understanding of the cell states found in these features, we applied our deconvolution method to the transcriptional profiles from the microdissected features of 10 patients and found they each exhibited a distinct cell state composition profile (**Figure 2A**). Leading-edge samples have been shown to exhibit expression patterns associated with the proneural subtype as well as neural tissue, suggesting they are composed of a mixture of tumor and normal cells (Gill et al., 2014; Jin et al., 2017; Puchalski et al., 2018). Consistent with this finding, we found this region was rich in oligodendrocytes found at the tumor-normal brain interface and was also predicted to contain high levels of stem-like malignant cells, despite its reduced tumor content. We have previously shown that stem-like cells and a subset of differentiated-like cells resemble a malignant oligodendroglial precursor cell-like state that has been implicated in neuronal signaling and synapse formation, suggesting transcriptional overlap between neural and tumor tissue in this region (Johnson et al., 2020; Venkatesh et al., 2019). Pseudopalisading cells around necrosis features, which are areas of hypoxia, exhibited the highest levels of differentiated-like malignant cells. Conversely, microvascular proliferation features were enriched in proliferating stem-like malignant cells, supporting the role of oxygen in influencing cell state. Finally, the cellular tumor feature exhibited more sample-specific variation, with high levels of differentiated-like malignant cells in IDH-wild-type samples and high levels of stem-like cells in IDH-mutant samples. Each cell state’s distribution was more significantly associated with the histological feature than the patient from which it was derived (two-way ANOVA; **Figure S2A**) (Puchalski et al., 2018).

**Figure 2.**
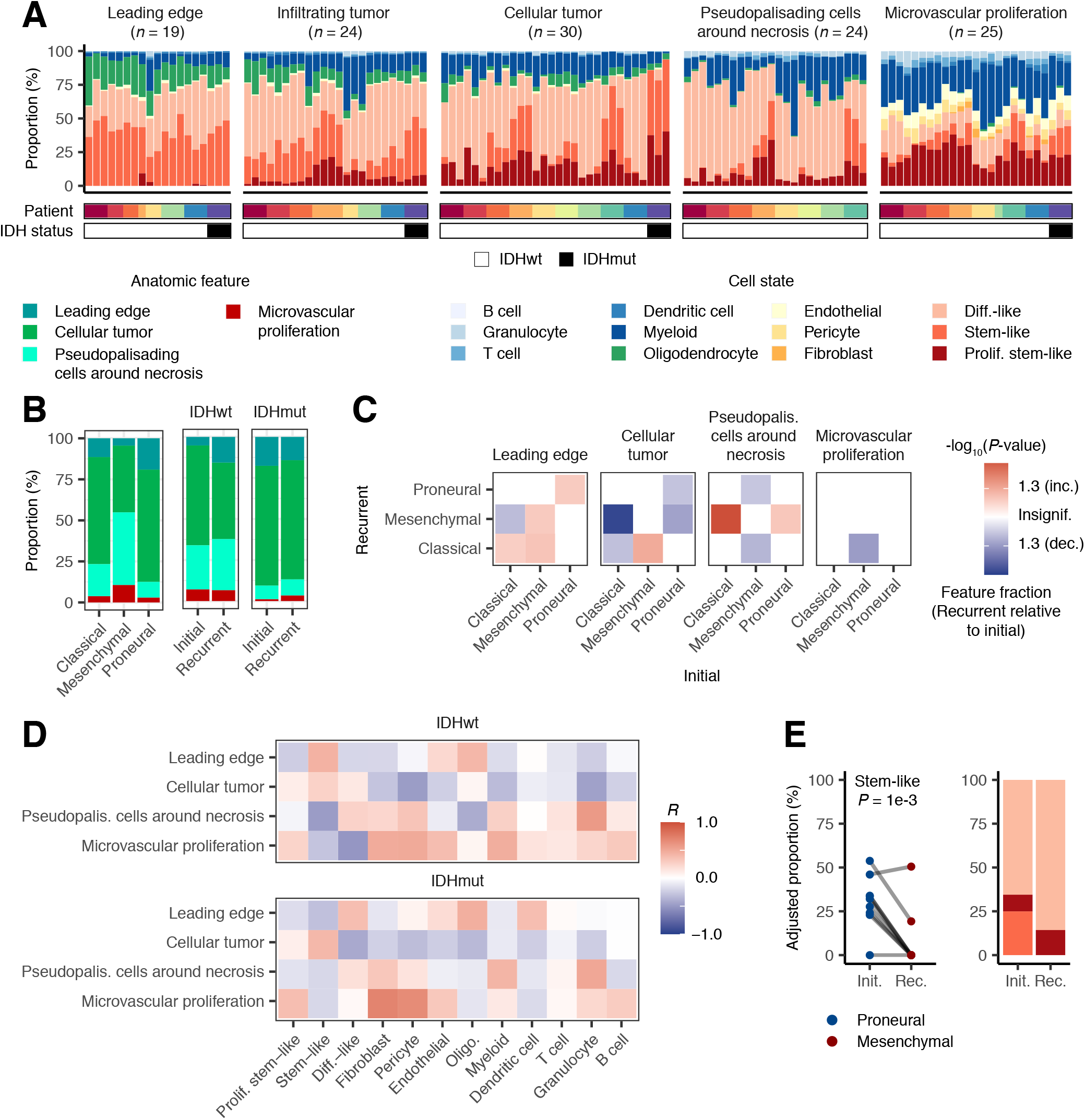
Histological features underlie changes in the cellular composition of diffuse glioma over time. (A) The cell state composition of each of the reference histology-defined Ivy GAP histological features from 10 patients. Patient and IDH mutation status tracks are included beneath the stacked bar plots. For the patient track, each colored block represents a unique patient. (B) Left: The average histological feature composition of each bulk transcriptional subtype for all initial GLASS tumors. Right: The average histological feature composition of initial and recurrent tumors stratified by IDH mutation status. (C) Heatmap depicting the significance of the changes in each histological feature between initial and recurrent tumors undergoing the indicated subtype transition. The initial subtype is indicated in the columns and the recurrent subtype is indicated in the rows. Colors represent the -log_10_(*P*-value) from a paired t-test, with increases at recurrence colored in red, decreases colored in blue, and *P*-values > 0.05 colored white. (D) Heatmap depicting the Pearson correlation coefficients measuring the association between the change in a given histological feature and the change in a given cell state when going from an initial tumor to recurrence. (E) Left: Ladder plot depicting the change in the adjusted stem-like cell proportion between paired initial and recurrent tumors undergoing a proneural-to-mesenchymal transition. Right: The average adjusted proportions for malignant cells for the tumor pairs outlined on the left. Malignant cell proportions were adjusted for the presence of non-malignant cells as well as non-cellular tumor content.

Given the strong association between histological features and cellular composition, we examined how the representation of these features varied over time by deconvoluting the GLASS dataset with the available feature-specific gene signatures developed as part of Ivy GAP. This analysis captured differences in each bulk transcriptional subtype’s anatomy that reflected their underlying cell state composition (**Figure 2B**). It also revealed that IDH-wild-type tumors had significantly higher leading-edge content at recurrence, even after adjusting for transcriptional subtype switch, which was consistent with the increase in oligodendrocytes we had previously observed (*P* = 1e- 4, paired t-test; **Figures 2B, 2C**). In IDH-wild-type tumors undergoing the common classical-to-mesenchymal transition, we observed a significant increase in pseudopalisading cells around necrosis and a decrease in cellular tumor content, indicative of increased hypoxia and non-malignant content (*P* = 2e-5, and 3e-5, respectively, paired t-test). At the cell state level, we found that changes in the abundance of differentiated-like malignant cells positively associated with increased cellular tumor features in IDH-wild-type tumors, increased leading edge features in IDH-mutant tumors, and increased pseudopalisading cells around necrosis features in both subtypes. Changes in stem-like malignant cells positively associated with changes in leading-edge features in IDH-wild-type tumors and cellular tumor features in IDH-mutant tumors. Finally, in both subtypes, changes in proliferating stem-like and immune cells positively associated with changes in microvascular proliferation (**Figure 2D**).

Given these correlations, we hypothesized that subtype switches in IDH-wild-type tumors were attributable to changes in histological feature composition over time. We recalculated our malignant cell fractions by adjusting for the presence of non-malignant cells, as well as leading-edge content which may vary by surgery. While most subtype switches associated with changes in at least one malignant cell fraction pre-adjustment, the only difference observed post-adjustment was a decrease in stem-like cells in tumors undergoing a proneural-to-mesenchymal transition (*P* = 3e-4, paired t-test; **Figures S2B, S2C**). These associations remained significant even after adjusting for the remaining non-cellular tumor features, suggesting tumors undergoing this switch exhibit a loss of stem-like cells independent of histological feature composition (**Figures 2E, S2B**). Collectively, these results indicate that while most subtype switches in IDH-wild-type tumors are related to changes in a tumor’s underlying physical structure and microenvironment, the changes observed in the proneural-to-mesenchymal transition may result from tumor-wide changes that reflect malignant cell-intrinsic processes at recurrence.

### Acquired somatic alterations at recurrence associate with changes in cellular composition

Somatic genetic alterations have been shown to be associated with the cell state distribution of IDH-wild-type and IDH-mutant glioma (Neftel et al., 2019; Tirosh et al., 2016; Verhaak et al., 2010). We thus hypothesized that changes in cellular composition resulted from genetic changes at recurrence. This was reinforced by the observation that, in both IDH-wild-type and IDH-mutant tumors, each cell state’s initial fractions weakly correlated with those at recurrence (median concordance coefficient (ρ_C_) = 0.17 and 0.26, respectively; **Figure 3A**). We reasoned that if the presence of a cell state was influenced by genetic factors, the pairwise change in its proportion over time would deviate from a zero-centered normal distribution that is suggestive of stochastic change.

**Figure 3.**
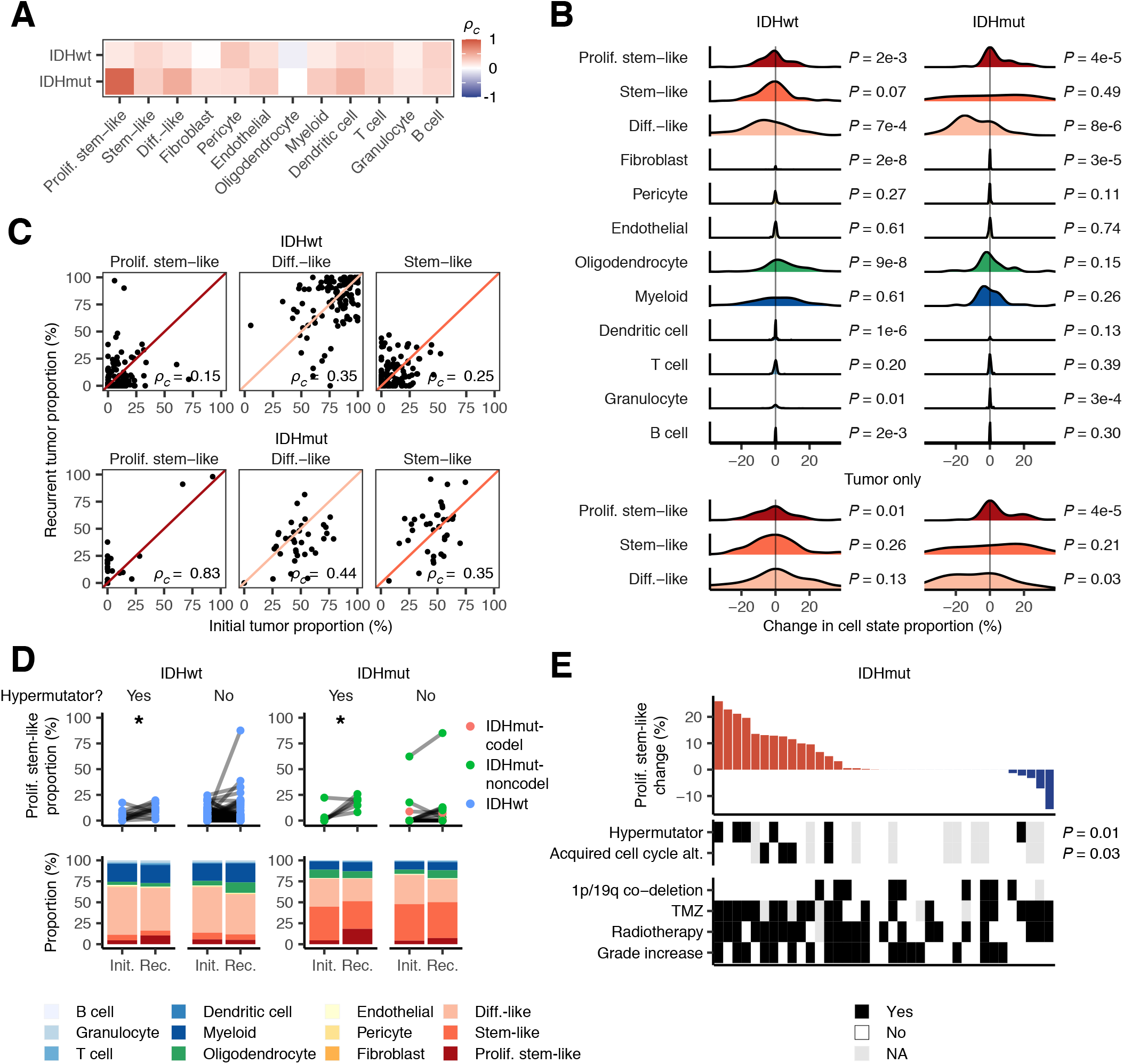
Hypermutation and acquired cell cycle alterations associate with increased proliferating stem-like malignant cells in IDH-wild-type and IDH-mutant glioma. (A) Heatmap depicting the concordance coefficients measuring the association between the indicated cell state fractions between initial and recurrent tumors. (B) Top: Density plots depicting the cell state proportion change distribution for each of the indicated cell states. Samples are stratified based on IDH mutation status. The tumor-only distributions indicate the change in malignant cell fractions after adjusting for non-malignant cells. *P*-values were derived using the Kolmogorov- Smirnov test that compared each distribution to a normal distribution with a mean of 0. (C) Scatterplots depicting the association between the adjusted malignant cell proportions in initial and recurrent tumors. Concordance coefficients are indicated. Diagonal lines correspond to the line y = x. (D) Top: Ladder plots depicting the change in the proliferating stem-like cell proportion between paired initial and recurrent tumors that did and did not undergo hypermutation. Point colors indicate IDH mutation and 1p/19q co-deletion status. * indicates paired t-test *P*-value < 0.05. Bottom: The average proportions of each cell state for the tumor pairs outlined above. (E) Top: The change in proliferating stem-like cell fraction between initial and recurrent tumors from IDH-mutant pairs. Each bar represents a tumor pair. Bottom: Molecular and clinical information for each tumor pair. *P*-values were calculated using a paired t-test measuring the association between initial and recurrent tumors that acquired the indicated phenotypes.

When we examined the distribution of each malignant cell state’s changes, we found that proliferating stem-like malignant cells significantly deviated from the stochastic distribution in IDH-mutant and IDH-wild-type glioma, and this remained true after adjusting for the presence of non-malignant cells (*P* < 0.05, Kolmogorov-Smirnov test; **Figure 3B, 3C**). Notably, we did not observe a change in stem-like cells, though we did not adjust for histological feature composition as we were focused on tumor-wide changes in cell state composition. Within IDH-mutant tumors, we identified acquired deletions of the cell cycle regulator *CDKN2A* and acquired amplifications of the cell cycle regulator *CCND2* as genetic events that together associated with the increase in proliferating stem-like cells (*P* = 0.01, paired t-test, *n* = 3; **Figure S3A**). This association was not present in IDH-wild-type tumors, which typically harbor *CDKN2A* deletions at initial presentation. Approximately 20% of gliomas recur with a hypermutated phenotype following treatment with alkylating agents, a standard-of-care chemotherapy (Barthel et al., 2019; Touat et al., 2020). This phenotype has been associated with disease progression and distant recurrence (Yu et al., 2021). We found that in both IDH-wild-type and IDH-mutant glioma, hypermutation also associated with an increase in proliferating stem-like malignant cells (*n* = 12 and 6, respectively; **Figure 3D**). In IDH-mutant tumors, hypermutation was independent of acquired copy number changes in *CDKN2A* and *CCND2*, suggesting that there are multiple genetic routes to increasing proliferating stem-like malignant cells at recurrence (**Figure 3E**). Notably, we found that neither hypermutation nor acquired cell cycle alterations were associated with changes in microvascular proliferation, suggesting that the increase in proliferating stem-like malignant cells in these tumors was driven by changes in their genetics (**Figure S3B**).

Beyond malignant cells, we observed that fibroblasts, oligodendrocytes, and granulocytes all deviated from the stochastic distribution. As with the proliferating stem-like cells, we compared how each cell state fraction differed in the small number of samples that acquired or lost selected driver mutations at recurrence. In IDH-wild-type tumors, tumors acquiring *NF1* mutations all underwent a mesenchymal transition and exhibited a significant increase in granulocytes (*P* = 0.01, paired t-test, *n* = 6; **Figure S3C**). Granulocytes have previously been associated with tumor necrosis, a feature that is prominent in mesenchymal glioblastoma (Yee et al., 2020). There were additionally several copy number alterations, including loss of *EGFR* or *PDGFRA* amplifications, that were associated with increased non-malignant cell content (*P* < 0.05, paired t-test, *n* = 9 and *n* = 3, respectively), and a transition to the mesenchymal subtype (*P* = 0.02, Fisher’s exact test; **Figures S3D and S3E**). We did not observe any significant changes in the fractions of non-malignant cells when comparing hypermutated recurrences with their corresponding non-hypermutated initial tumors, although T cells numerically increased in IDH-mutant tumors (*P* = 0.07, paired t-test; **Figure S3F**). Collectively, nearly all the cell state changes we found to deviate from the stochastic distribution were associated with changes in tumor genetics, suggesting that genetic evolution underlies the most frequent changes in cellular composition over time.

### IDH-wild-type malignant cells exhibit an increase in neuronal signaling gene expression programs at recurrence

While a subset of tumors demonstrated increases in proliferating stem cell content at recurrence, most IDH-wild-type and IDH-mutant tumors did not exhibit any directed changes in their malignant cell composition over time. We hypothesized that the expression programs of individual cell states may change following treatment in more subtle ways that do not manifest as a noticeable shift in cellular composition. To test whether these changes were taking place, we utilized our pan-glioma single-cell RNAseq dataset as a reference to deconvolute GLASS bulk gene expression profiles into their component differentiated-like, stem-like, proliferating stem-like, and myeloid gene expression profiles (**Figure S4A**). Comparing these profiles to those derived from fluorescence-activated cell sorting (FACS)-purified glioma-specific CD45^-^ and myeloid populations revealed strong concordance between the corresponding profiles of each cell state (**Figures S4B and S4C**).

To compare how the expression programs in each malignant cell state vary longitudinally, we compared the cell state-specific gene expression profiles between the initial and recurrent tumor for each pair receiving temozolomide and/or radiotherapy. We only included tumor pairs that did not exhibit a bulk transcriptional subtype switch, as variable subtype switching may reflect changes in histological feature composition over time. In IDH-wild-type tumors, we found that 5.2% of the 7,400 genes that could be inferred in stem-like cells were significantly differentially expressed at recurrence (false discovery rate (FDR) < 0.1, Wilcoxon signed-rank test). This number was 1.9% of the 11,376 differentiated-like state genes and 0.5% of the 5,908 proliferating stem-like state genes (**Figure 4A**; **Table S4**). Based on these results, we defined recurrence-specific signatures as the genes that were significantly up-regulated at recurrence in each cell state. While there was little overlap between each of these signatures, gene ontology (GO) enrichment analysis revealed that the stem-like and differentiated-like signatures were significantly enriched in terms relating to neuronal signaling (**Figures 4B and S4D**). These results were consistent with the increase in leading edge features and oligodendrocytes at recurrence we had previously observed. To confirm that these signatures were measuring malignant-specific expression changes at recurrence, we compared how their expression differed between malignant single cells from unmatched initial and recurrent IDH-wild-type tumors, as there is limited availability of matched initial and recurrent single-cell data. In all cases, the recurrence-specific cells exhibited significantly higher expression of their respective signatures than those from initial tumors (**Figure 4C**). We next examined each recurrence-specific signature’s association with histological feature content and found that the tumor’s leading edge was positively associated with the malignant cell state-specific expression of each signature (**Figure 4D**). While this feature has reduced tumor content, malignant cells in the tumor periphery have previously been shown to exhibit neuronal signaling activity (Darmanis et al., 2017; Puchalski et al., 2018). Furthermore, stem-like cells, which are the malignant state most frequently found at the leading edge and enhancing region (Jin et al., 2017), exhibited the strongest associations. Notably, each of these associations was present regardless of whether the comparisons were made in initial or recurrent tumors. Together these results suggest that increased normal cell content at recurrence associates with higher signaling between malignant cells and neighboring neural cells. Neuron-to-glioma synapses have been implicated in increased tumor growth and invasion, and collectively our results support a model of greater tumor invasion into the normal brain at recurrence that is facilitated by an increase in neuronal interactions (Venkataramani et al., 2019; Venkatesh et al., 2015; Venkatesh et al., 2019; Venkatesh et al., 2017).

**Figure 4.**
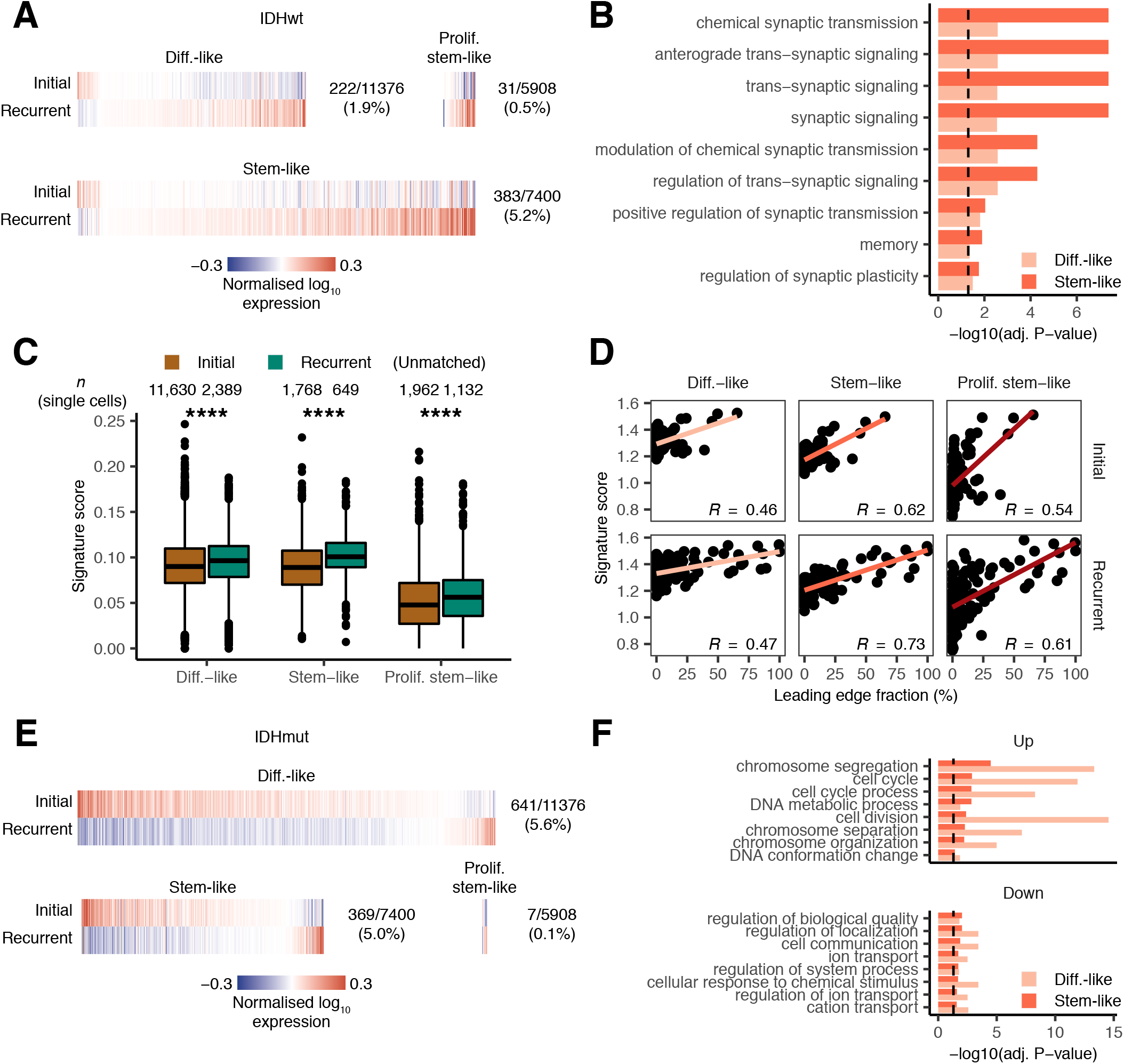
Malignant cells exhibit increased neuronal signaling and cell cycle activation programs in recurrent IDH-wild-type and IDH-mutant tumors. (A) Heatmaps depicting the average normalized log_10_ expression level of genes that were differentially expressed between malignant cell states from initial and recurrent IDH-wild-type tumors not undergoing a subtype switch. Fractions on each plot’s right indicate the number of differentially expressed genes (numerator) out of the number of genes inferred for that cell state’s profile using CIBERSORTx (denominator). (B) Bar plot depicting the -log_10_(adjusted *P*-value) from a GO enrichment analysis for the differentially expressed genes in differentiated-like and stem-like malignant cells depicted in (A). Only GO terms that were enriched at an adjusted *P*-value of < 0.05 in both the differentiated-like and stem-like signatures were included. (C) Boxplot depicting the average signature expression in single cells of the indicated malignant cell states from unmatched initial and recurrent IDH-wild-type tumors. **** indicates Wilcoxon rank-sum test *P*-value < 1e-5. (D) Scatterplot depicting the association between the leading edge fraction and the average signature expression in the inferred malignant cell state-specific expression profiles of samples in the GLASS dataset. Pearson correlation coefficients are indicated. (E) Heatmaps depicting the average normalized log_10_ expression level of genes that were differentially expressed between malignant cell states from initial and recurrent IDH-mutant tumors not undergoing a subtype switch. Fractions are as outlined in (A). (F) Bar plots depicting the -log_10_(adjusted *P*-value) from a GO enrichment analysis for the differentially expressed genes in differentiated-like and stem-like malignant cells depicted in (E). Top 8 GO terms that were significant in the up- or down-regulated signatures from differentiated-like and stem-like cells are shown. In (B) and (F), dotted line corresponds to adjusted *P*-value < 0.05.

We next compared how the expression profiles of each cell state differed between initial and recurrent IDH-mutant tumors that received treatment. The resulting signatures were distinct from those in IDH-wild-type tumors, with the largest number of differentially expressed genes found in the differentiated-like state instead of the stem-like state (FDR < 0.1, Wilcoxon signed-rank test; **Figure 4E**, **Table S4**). Additionally, the majority of candidate genes identified in IDH-mutant tumors were expressed more highly in initial tumors, as opposed to IDH-wild-type tumors where the reverse was true. As with IDH-wild-type tumors, there was limited overlap between the differentiated-like and stem-like signatures (**Figure S4E**). A GO enrichment analysis of the genes up-regulated at recurrence in the differentiated-like and stem-like cell states revealed an enrichment of cell cycle-related genes. In contrast, the down-regulated genes were enriched in terms related to cellular communication and response to stimulus (**Figure 4F**). These signatures were consistent with those found in higher grade tumors, suggesting that the cell state-specific gene expression changes were indicative of grade increases at recurrence. Accordingly, we observed that these changes were strongest in the tumor pairs that recurred at a higher grade (**Figure S4F**). Furthermore, when we compared signature expression in single cells of the same cell state, we found that the signatures were differentially expressed in the cells derived from grade III versus grade II tumors (**Figure S4G**). These results indicate that IDH-wild-type and IDH-mutant tumors recur in distinct manners that may reflect their response to treatment.

### Mesenchymal tumor cell activity associates with a distinct myeloid cell phenotype

The mesenchymal subtype of glioma is associated with increased accumulation of immune cells, primarily of the myeloid lineage (Bhat et al., 2013; Kim et al., 2021; Wang et al., 2017). We thus hypothesized that interactions between the tumor-infiltrating myeloid cells and malignant cells can influence the tumor’s trajectory at recurrence. To understand how the myeloid compartment differed across each glioma subtype, we deconvoluted the myeloid-specific gene expression profiles from a collection of diffuse glioma bulk RNAseq profiles (*n* = 701) from The Cancer Genome Atlas (TCGA). The myeloid compartment in IDH-wild-type tumors was characterized by high expression of a previously defined blood-derived macrophage signature (Muller et al., 2017), while myeloid cells in IDH-mutant-noncodel tumors exhibited high expression of a previously defined brain-resident microglia signature (**Figure 5A**). Stratifying this cohort by transcriptional subtype revealed that the blood-derived macrophage signature followed a stepwise increase with mesenchymal subtype representation, while microglial gene expression was highest amongst tumors of the mixed subtype classification that is seen most frequently in IDH-mutant-noncodel glioma (**Figure S5A**). In IDH-wild-type tumors, blood-derived macrophage signature expression was positively correlated with the abundance of microvascular proliferation and pseudopalisading cells around necrosis features, while the microglia signature was most positively correlated with leading-edge content. There were no clear associations for either signature in IDH-mutant tumors (**Figure S5B**). Longitudinally, when holding transcriptional subtype constant, we observed very few differentially expressed genes in the myeloid cell profiles from matched initial and recurrent tumors in the GLASS cohort (**Figure S5C**). However, the myeloid profiles in IDH-mutant tumors that increased grade at recurrence exhibited a significant decrease in microglia signature expression, suggesting a shift in myeloid cell states away from brain-resident microglia (*P* = 1e- 3, Wilcoxon signed-rank test; **Figure 5B**).

**Figure 5.**
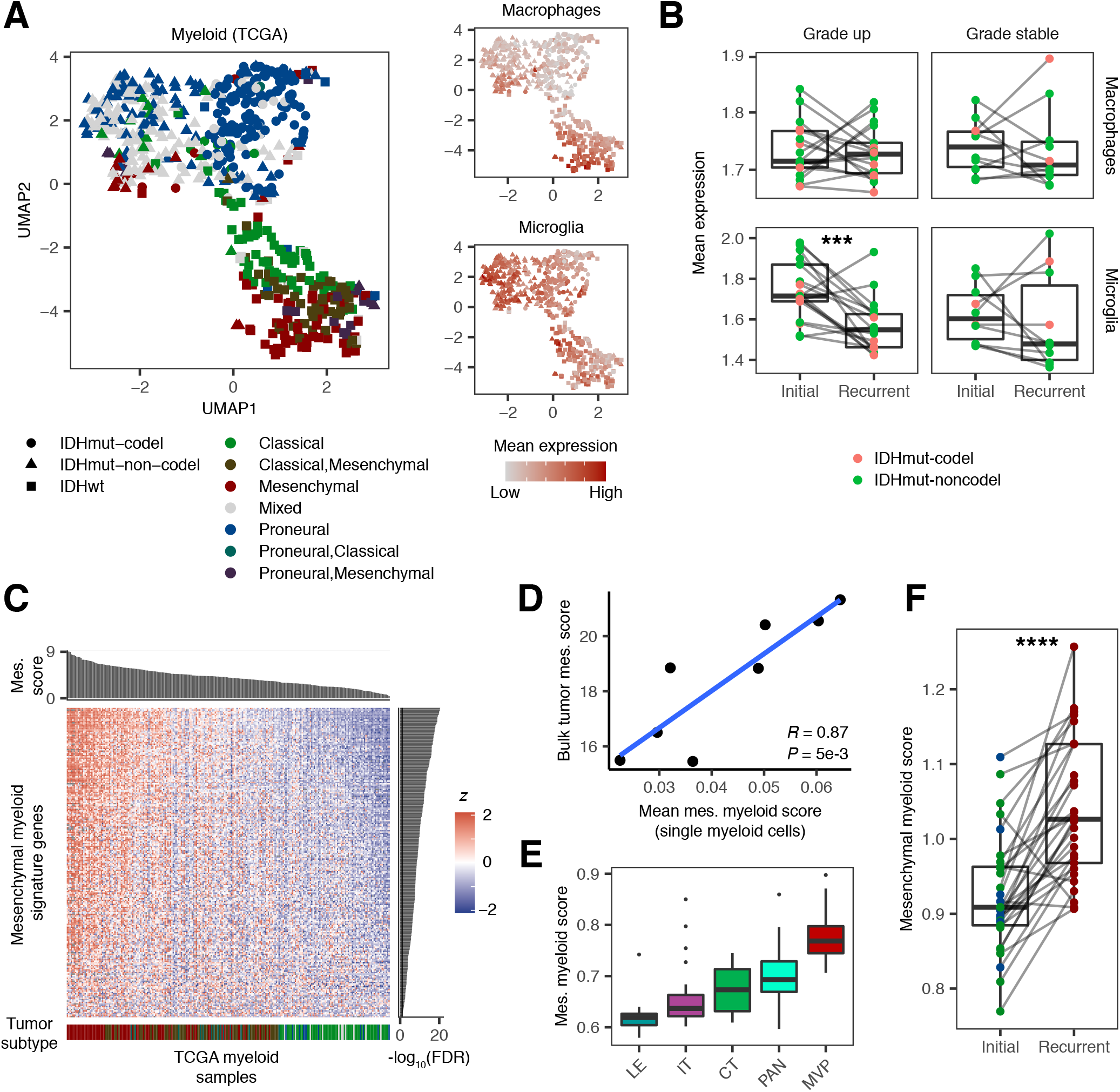
Myeloid cells in diffuse glioma exhibit diverse phenotypes based on IDH mutation status, transcriptional subtype, and recurrence status. (A) Left: Uniform Manifold Approximation and Projection (UMAP) dimensionality reduction plot of the CIBERSORTx-inferred myeloid profiles from TCGA. Colors indicate bulk transcriptional subtype; shapes indicate IDH and 1p/19q co-deletion status. When all three bulk transcriptional subtypes were significantly represented in a sample, the ‘mixed’ classification was used. Right: UMAP plot colored based on the relative mean expression of macrophage and microglia signatures (B) Box and ladder plots depicting the difference in the mean expression of the indicated signatures between initial and recurrent IDH-mutant tumors from GLASS that do and do not recur at higher grades. Point colors indicate 1p/19q co-deletion status. *** indicates Wilcoxon signed-rank test *P*-value < 1e-3. (C) Heatmap depicting the normalized expression z-score of genes that were differentially expressed between myeloid cells from mesenchymal and non-mesenchymal TCGA tumors. Rows indicate genes and columns indicate samples. Top sidebar indicates the bulk mesenchymal score of each sample divided by 1,000. Right sidebar indicates the -log_10_ adjusted Wilcoxon rank-sum test *P*- value of the association for each gene. Bottom sidebar indicates the transcriptional subtype of each sample per panel (A). (D) Scatterplot depicting the association between the mean mesenchymal myeloid signature expression in single myeloid cells and the mesenchymal subtype score calculated from bulk RNAseq for each patient. (E) Boxplot depicting the mean mesenchymal myeloid signature expression for CIBERSORTx-inferred myeloid profiles from different histological features in the Ivy GAP dataset. Features in this dataset include the leading edge (LE), infiltrating tumor (IT), cellular tumor (CT), pseudopalisading cells around necrosis (PAN), and microvascular proliferation (MVP). (F) Box and ladder plots depicting the difference in the mean expression of the mesenchymal myeloid signature between initial and recurrent IDH-wild-type tumors undergoing a mesenchymal transition in GLASS. **** indicates Wilcoxon signed- rank test *P* < 1e-5.

Macrophages are highly plastic and capable of changing their transcriptional programs in response to different stimuli (Xue et al., 2014). We reasoned that interactions between myeloid cells and malignant cells in the mesenchymal glioma microenvironment might result in a population of myeloid cells that bear a distinct transcriptional phenotype. We thus performed a differential expression analysis to compare how the deconvoluted myeloid cell expression profiles differed between mesenchymal and non-mesenchymal IDH-wild-type tumors in TCGA. This analysis revealed that 218 of the 4,235 inferred genes (5%) were significantly upregulated in mesenchymal samples (FDR < 0.1, fold-change > 1.1; **Figure 5C**, **Table S5**). When we examined the average expression of this signature in myeloid cells from our scRNAseq dataset, we found that the average signature score in each patient was strongly associated with the mesenchymal glioma subtype score derived from their patients’ respective bulk RNAseq profile (*R* = 0.87, *P =* 5e-4; **Figure 5D**). We did not observe this association using the blood-derived macrophage signature, suggesting that our mesenchymal macrophage signature was measuring myeloid activity specific to the mesenchymal subtype (**Figure S5D**). Analysis of signature expression across each of the Ivy GAP dataset’s histological feature samples revealed that the mesenchymal myeloid signature was expressed most highly in the pseudopalisading cells around necrosis and microvascular proliferation features that are highest in mesenchymal tumors (**Figure 5E**). A GO enrichment analysis of this signature revealed the mesenchymal myeloid signature to be enriched in chemokine signaling and lymphocyte chemotaxis functions (**Figure S5E**).

Longitudinally, IDH-wild-type tumors in the GLASS dataset undergoing a mesenchymal transition at recurrence exhibited significantly higher mesenchymal myeloid signature expression in their recurrent tumor myeloid profiles (*P* = 6e-7, Wilcoxon signed-rank test; **Figure 5F**). This led us to examine whether we could identify the ligand-receptor interactions between myeloid and malignant cells associated with this transition over time. We focused this analysis on differentiated-like malignant cells, as this cell state frequently exhibits mesenchymal-like characteristics (Johnson et al., 2020). To probe these interactions, we downloaded a set of 1,894 literature-supported ligand-receptor pairs (Ramilowski et al., 2015) and identified all pairs that had one component expressed in a tumor’s deconvoluted myeloid profile and the other expressed in the differentiated-like malignant cell profile. We then compared how the longitudinal change in expression of each component associated with the change in each tumor pair’s mesenchymal subtype score. This identified 69 putative ligand-receptor pairs where each component exhibited a positive association (*R* > 0, FDR < 0.1; **Figure S5F**). Of these pairs, 35 also exhibited these associations in our single-cell dataset, including 19 where the ligand was expressed by the malignant cell and 16 where the ligand was expressed by the myeloid cell (**Table S6**). In pairs where the ligand was expressed by the malignant cell, the pair with the highest mean correlation was vascular endothelial growth factor A (*VEGFA*)-neuropilin 1 (*NRP1*), which is involved in angiogenesis and endothelial cell migration (Herzog et al., 2011). In pairs where the myeloid cell expressed the ligand, the best performing pair was oncostatin M (*OSM*)-oncostatin M receptor (*OSMR*), which has been associated with an epithelial-to-mesenchymal transition *in vitro* (Junk et al., 2017). In addition to these pairs, myeloid-specific single-cell expression of the receptor *MARCO* was significantly associated with the bulk tumor mesenchymal signature score, in concordance with its reported role as a marker of mesenchymal-associated macrophages (**Figure S5G**) (Sa et al., 2020). These analyses identify candidate receptor-ligand interactions that can potentially be targeted to shift a tumor towards or away from a mesenchymal state following treatment.

### Antigen presentation is disrupted at recurrence in IDH-mutant-noncodel glioma

Studies in non-small cell lung cancer and other cancer types have shown that cytotoxic T cells exert selective pressure on malignant cells through the elimination of neoantigen-presenting tumor subclones (Grasso et al., 2018; McGranahan et al., 2017; Rooney et al., 2015; Rosenthal et al., 2019; Zhang et al., 2018). Immune interactions have been associated with selection for epigenetic changes in glioma (Gangoso et al., 2021), however the extent to which T cells are involved in shaping genetic evolution of glioma remains unclear. We hypothesized that if T cell selection was taking place, then tumors with high T cell infiltration would more frequently exhibit loss-of-heterozygosity (LOH) in the human leukocyte antigen (HLA) genes that are central to the presentation of neoantigens. We thus called HLA LOH throughout the GLASS cohort (**Figure 6A**). We observed that HLA LOH is prevalent in glioma, occurring in at least one timepoint in 19% of patients. Within IDH-wild-type and IDH-mutant-codel tumors, HLA LOH was found at similar rates between initial and recurrent tumors, with most affected pairs exhibiting this alteration at both time points. This was not the case in IDH-mutant-noncodel tumors, where significantly more samples acquired HLA LOH at recurrence (*P* = 0.02, Fisher’s exact test). However, unlike in non-small cell lung cancer, the presence of HLA LOH was not associated with the fraction of infiltrating T cells in each tumor (**Figure 6B**). Furthermore, we did not observe an association between T cell abundance and the rates of neoantigen depletion, and in HLA LOH samples, the number of neoantigens binding to the kept allele did not differ from the number that were predicted to bind to the lost allele (**Figures S6A and S6B**).

**Figure 6.**
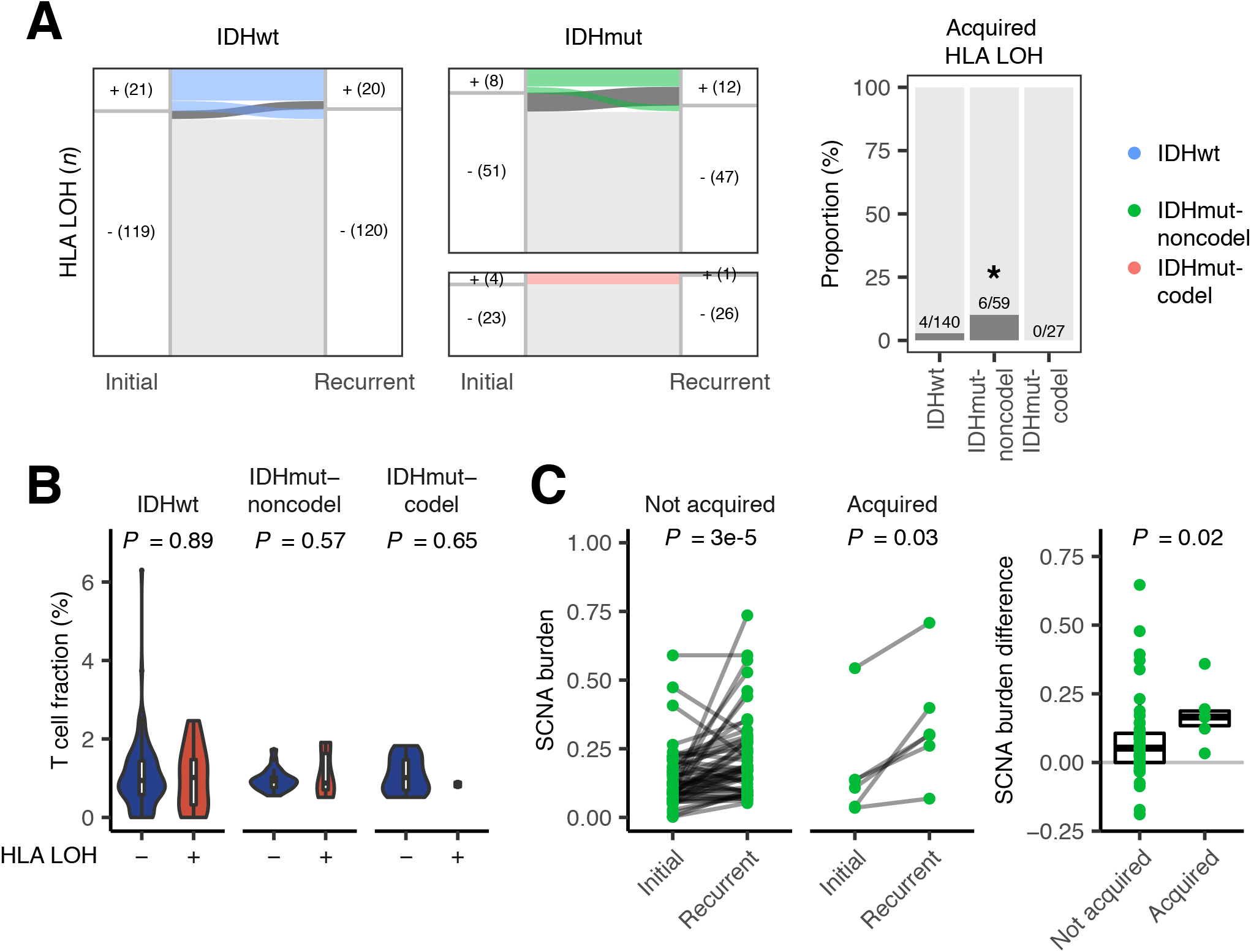
Loss of heterozygosity in HLA genes is associated with increased somatic copy number alterations in IDH-mutant non-1p/19q co-deleted glioma. (A) Left: Sankey plot indicating whether a tumor pair acquires or loses HLA LOH at recurrence. Colored lines reflect the IDH and 1p/19q co-deletion status of the tumor pair and indicate HLA LOH in the initial tumor. Dark gray lines indicate acquired HLA LOH. Right: Stacked bar plot indicating the proportion of samples of each glioma subtype that acquired HLA LOH at recurrence. * indicates Fisher’s exact test *P*-value < 0.05. (B) Violin plot depicting the difference in T cell proportion in samples with and without HLA LOH. *P*-values were calculated using the t-test. (C) Left: Ladder plots depicting the change in SCNA burden between paired initial and recurrent IDH-mutant-noncodel tumors that did and did not acquire HLA LOH. *P*-values were calculated using the Wilcoxon signed-rank test. Right: Boxplot depicting the difference in the change in SCNA burden between IDH-mutant- noncodel tumor pairs that did and did not acquire HLA LOH. *P*-value was calculated using the Wilcoxon rank-sum test.

Given the absence of an association between HLA LOH status and T cell infiltration, we reasoned that HLA LOH might be a passenger event that occurs in samples with a high genome-wide somatic copy number alteration (SCNA) burden. We had previously shown that IDH-mutant- noncodel tumors exhibit significantly higher SCNA burdens at recurrence (Barthel et al., 2019). This difference remained significant regardless of whether the tumors acquired HLA LOH. However, the tumors acquiring this alteration at recurrence exhibited significantly higher changes in SCNA burden than those that did not, confirming our hypothesis (*P* = 0.02, Wilcoxon rank-sum test; **Figure 6C**). We did not observe longitudinal associations between HLA LOH status and SCNA burden in IDH-wild-type tumors, although we found at both the initial and recurrent time points that samples with HLA LOH had higher SCNA burdens than those with both HLA alleles (**Figure S6C**). Taken together, these results suggest that disruption of antigen presentation in glioma is likely a byproduct of SCNA burden rather than being a result of selection by cytolytic T cells as has been observed in other cancers.

## Discussion

To understand the factors driving the evolution and treatment resistance of diffuse glioma, we integrated genomic and transcriptomic data from the initial and recurrent tumor pairs of 292 patients. By integrating this resource with data from single-cell RNAseq experiments, a histological transcriptional atlas, and a multitude of external transcriptional datasets, we have comprehensively defined the longitudinal transcriptional and compositional changes that gliomas sustain at recurrence.

In this study, we employed single-cell deconvolution approaches to enable high-resolution quantification of glioma tumors’ cellular composition. Available cell state classification models have been developed for diffuse glioma using single cells of a single glioma subtype (Castellan et al., 2021; Garofano et al., 2021; Neftel et al., 2019; Richards et al., 2021; Venteicher et al., 2017). In contrast, our reference matrix utilized cell states derived from a pan-glioma single-cell dataset composed of initial and recurrent tumors of all major clinically relevant glioma subtypes, and thus included malignant and normal cell states commonly found across diffuse glioma. The resulting cellular proportions reflected true cell state levels in multiple benchmarking analyses, making this an invaluable approach for comparing and contrasting the longitudinal changes taking place across IDH-wild-type and IDH-mutant tumors. In the future this approach can continue to be refined as the number of cells per tumor and patients profiled by scRNAseq increases and enables even higher resolution estimates of glioma cell state composition and heterogeneity.

While transcriptional subtype switching has been reported to occur frequently in IDH-wild-type glioma, the role these switches play in treatment resistance is unclear. Pathology-defined histological features from Ivy GAP exhibit distinct transcriptional profiles that correspond to different glioma transcriptional subtypes, suggesting that subtype switching may be more reflective of changes in the tumor’s histological feature composition at recurrence (Jin et al., 2017; Puchalski et al., 2018). Ivy GAP comprises features defined from primary tumors, which we found to be useful proxies to measure the biological changes at recurrence that underlie subtype switching. Limitations of the Ivy GAP resource may include the absence of commonly observed features, such as necrotic tissue and perinecrotic zone tumor, which may be more present following radiation therapy. We showed that the proneural-to-mesenchymal transition is independent of histological feature composition and reflects transcriptional changes in the cellular tumor. Mesenchymal transitions have been shown to associate with several factors, including increased myeloid cell infiltration, radiation-induced NF-κB activation, altered tumor metabolism, and hypoxia (Bhat et al., 2013; Garofano et al., 2021; Kim et al., 2021; Mao et al., 2013; Osuka et al., 2021; Schmitt et al., 2021; Wang et al., 2017). Our results indicate that the proneural-to-mesenchymal transition is likely influenced by tumor-wide changes, supporting the hypothesis that this transition is involved in therapy resistance. Additional studies where multiple biopsies are obtained from the same tumor over time may help to further elucidate the relationship between histological feature composition and gene expression subtype.

Across IDH-wild-type and IDH-mutant glioma, we identified a sub-population of samples that exhibited an increase in proliferating stem-like malignant cells at recurrence. Analysis of the acquired somatic alterations in these tumors revealed that hypermutation was associated with this change in both subtypes. This finding across both subtypes suggests that hypermutation may represent a pan-glioma treatment resistance mechanism. Hypermutation did not associate with patient survival in the GLASS dataset but has been found more frequently in distant recurrences and linked to reduced survival following high-grade progression in low-grade IDH-mutant tumors (Barthel et al., 2019; Touat et al., 2020; Yu et al., 2021). Given these findings, our data highlights methods to predict treatment-induced hypermutation represent a previously unrecognized unmet clinical need in the field. Integrating such methodologies into clinical care pathways would help to identify patients that may benefit from therapies that complement chemotherapy and further target cycling cells.

We did not identify any somatic alterations associated with changes in malignant cell composition outside of hypermutation and copy number changes in cell cycle regulators. Despite this, we found that malignant glioma cells in IDH-wild-type tumors exhibited a significant increase in the expression of genes involved in neuronal signaling. This change coincided with an increase in oligodendrocytes at recurrence that was independent of the extent of tumor resection, providing a medium for increased interactions between malignant and normal cells in the brain. Additionally, neuronal signaling was most significantly up-regulated within the malignant stem-like cells, which are found at the highest levels at the leading edge of the tumor and frequently resemble oligodendroglial precursor-like malignant cells involved in neuronal signaling (Venkatesh et al., 2019). Increased neuronal signaling has previously been reported in malignant cells that have infiltrated into the surrounding tissue in response to low oxygen content and our study extends these observations to glioma progression (Darmanis et al., 2017). Collectively these findings coupled with our results relating to proneural-to-mesenchymal transition support a model where recurrent IDH-wild-type tumors, in response to changes in hypoxia or tumor metabolism at recurrence, invade the surrounding peripheral tissue where they actively interact with neighboring neuronal cells. Given the growing appreciation of the role neuron-glioma interactions play in glioma invasion and progression, it will be critical to understand the extent to which these interactions facilitate tumor regrowth and treatment resistance (Venkataramani et al., 2019; Venkatesh et al., 2015; Venkatesh et al., 2019; Venkatesh et al., 2017).

In agreement with other studies, we found that the myeloid cell phenotype varied in relation to tumor subtype and malignant cell state (Klemm et al., 2020; Muller et al., 2017; Ochocka et al., 2021; Pombo Antunes et al., 2021; Venteicher et al., 2017). Notably, we found that this variation was most apparent in mesenchymal tumors, where myeloid cells exhibited a distinct transcriptional program. Ligand-receptor analyses revealed several candidate interactions involved in driving malignant and myeloid cells toward this mesenchymal phenotype. Resolving the directionality of these interactions, or determining whether additional factors mediate them, will be an important step toward understanding the contribution myeloid cells make in mesenchymal transformation. We did not observe any differences in T cell activity, nor did we observe evidence of T cell-mediated selection, making glioma distinct from several other cancers (Grasso et al., 2018; McGranahan et al., 2017; Rooney et al., 2015; Rosenthal et al., 2019; Zhang et al., 2018). Despite this, we did observe that antigen presentation in IDH-mutant-noncodel tumors is frequently disrupted at recurrence and is associated with increases in SCNA burden. These results should inform the design of T cell-based immunotherapies going forward, as standard-of-care therapies may inadvertently disrupt malignant cells’ ability to present neoantigens to T cells.

Therapy resistance remains a significant obstacle for patients with diffuse glioma and must be overcome to improve patient survival and quality of life. Overall, our results reveal that gliomas undergo changes in cell states that associate with changes in genetics and the microenvironment, providing a baseline towards building predictive models of treatment response. Taking into consideration the current histopathologic diagnostic criteria for gliomas and their longitudinal follow-up, future efforts by the GLASS Consortium are now underway. These include expansion of the cohort, integration of digitized tissue sections, and association with clinical and genomic datasets with radiographic imaging data (Bakas et al., 2020). Computational imaging studies have shown mounting evidence and promise in revealing imaging signatures associated with increased invasion and proliferation for glioma patients harboring particular mutations (Bakas et al., 2017; Binder et al., 2018; Fathi Kazerooni et al., 2020; Mang et al., 2020; Zwanenburg et al., 2020), and given their use in clinical monitoring, are highly complementary to the longitudinal datasets established here. Going forward, the transcriptional and compositional changes we have identified can be integrated with these imaging-based results to more broadly assess the molecular and microenvironmental heterogeneity of glioma and identify clinically targetable factors to aid in shaping a patient’s disease trajectory.

## The GLASS Consortium

Frederick S Varn^1^, Kevin C Johnson^1^, Taylor E Wade^1^, Tathiane M Malta^2^, Thais S Sabedot^3^, Floris P Barthel^1^, Hoon Kim^1^, Nazia Ahmed^4^, Indrani Datta^5^, Jill S Barnholtz-Sloan^6^, Spyridon Bakas^7,8^, Fulvio D’Angelo^9^, Hui K Gan^10^, Luciano Garofano^9^, Jason T Huse^11^, Mustafa Khasraw^12^, Emre Kocakavuk^1^, Simona Migliozzi^9^, D. Ryan Ormond^13^, Sun Ha Paek^14^, Erwin G Van Meir^15^, Annemiek M.E. Walenkamp^16^, Colin Watts^17^, Michael Weller^18^, Tobias Weiss^18^, Pieter Wesseling^19^, Kenneth Aldape^21^, Kristin D Alfaro^22^, Samirkumar B Amin^1^, Kevin J Anderson^1^, Christoph Bock^23,24^, Daniel J Brat^25^, Andrew Brodbelt^26,27^, Ketan R Bulsara^28^, Ana Valeria Castro^3^, Jennifer M Connelly^29^, Joseph F Costello^30^, John F de Groot^22^, Gaetano Finocchiaro^31^, Pim J French^32^, Peter V Gould^33^, Ann C Hau^34^, Chibo Hong^30^, Craig Horbinski^35^, Azzam Ismail^36^, Mathilde CM Kouwenhoven^37^, Anna Lasorella^38,39,40^, Peter S LaViolette^41^, Allison K Lowman^41^, Keith L Ligon^42^, Annette M Molinaro^30^, MacLean P Nasrallah^8^, Ho Keung Ng^43^, Simone P Niclou^34^, Johanna M Niers^19^, Joanna J Phillips^30^, Raul Rabadan^44^, Ganesh Rao^45^, Guido Reifenberger^46^, Nader Sanai^47^, Susan C Short^4^, Peter Sillevis Smitt^32^, Andrew E Sloan^48^, Marion Smits^49^, James M Snyder^3^, Hiromichi Suzuki^50^, Ghazaleh Tabatabai^51^, Georgette Tanner^4^, William H Tomaszewski^52^, Michael Wells^3^, Bart A Westerman^37^, Helen Wheeler^53^, Adelheid Woehrer^54^, Jichun Xie^55,56^, W.K. Alfred Yung^22^, Gelareh Zadeh^57^, Junfei Zhao^44^, Lucy F Stead^4^, Laila M Poisson^5^, Houtan Noushmehr^3^, Antonio Iavarone^9,38,39,40^, Roel GW Verhaak^1^

## Affiliations

1. The Jackson Laboratory for Genomic Medicine, Farmington, CT, USA
2. School of Pharmaceutical Sciences of Ribeirao Preto, University of São Paulo, Brazil. Ribeirao Preto, Sao Paulo, Brazil
3. Hermelin Brain Tumor Center, Henry Ford Health System, Detroit, MI, USA
4. University of Leeds, Leeds, UK.
5. Department of Public Health Sciences, Hermelin Brain Tumor Center, Henry Ford Health System, Detroit, MI, USA
6. Case Western Reserve University School of Medicine and University Hospitals of Cleveland, Cleveland, OH, USA
7. Department of Radiology, University of Pennsylvania, Philadelphia, PA, USA
8. Department of Pathology and Laboratory Medicine, University of Pennsylvania, Philadelphia, PA, USA
9. Institute for Cancer Genetics, Columbia University Medical Center, New York, NY, USA
10. Olivia Newton-John Cancer Research Institute, Austin Health, Melbourne, Australia
11. University of Texas MD Anderson Cancer Center, Houston, TX, USA
12. Preston Robert Tisch Brain Tumor Center at Duke, Department of Neurosurgery, Duke University Medical Center, Durham, North Carolina
13. Department of Neurosurgery, University of Colorado School of Medicine, Aurora, Colorado, USA
14. Seoul National University College of Medicine and Seoul National University Hospital, Seoul, Republic of Korea
15. Department of Neurosurgery, School of Medicine and O’Neal Comprehensive Cancer Center, University of Alabama at Birmingham, Birmingham, Alabama, USA.
16. Department of Medical Oncology, University Medical Center Groningen, Groningen, The Netherlands
17. Academic Department of Neurosurgery, Institute of Cancer and Genomic Sciences, University of Birmingham, Birmingham, UK
18. Department of Neurology, University Hospital Zurich, Zurich, Switzerland
19. Amsterdam University Medical Centers/VUmc, Amsterdam, The Netherlands
20. Princess Máxima Center for Pediatric Oncology, Utrecht, The Netherlands
21. National Cancer Institute, Bethesda, MD, USA 20842
22. Department of Neuro-Oncology, University of Texas MD Anderson Cancer Center, Houston, TX, USA
23. CeMM Research Center for Molecular Medicine of the Austrian Academy of Sciences, Vienna, Austria
24. Institute for Artificial Intelligence and Decision Support, CeMSIIS, Medical University of Vienna,Vienna, Austria
25. Department of Pathology, Northwestern University Feinberg School of Medicine, Chicago, IL, USA
26. The Walton Centre NHS Foundation Trust, Liverpool, UK
27. University of Liverpool, Liverpool, UK
28. Division of Neurosurgery, University of Connecticut/UConn Health, Farmington, CT, USA
29. Medical College of Wisconsin, Milwaukee WI, USA
30. Department of Neurosurgery, University of California San Francisco, San Francisco, CA, USA
31. Fondazione IRCCS Istituto Neurologico Besta, Milano, Italy
32. Department of Neurology, Erasmus University Medical Centre Rotterdam, Rotterdam, The Netherlands
33. service d’anatomopathologie, Hôpital de l’Enfant-Jésus du CHU de Québec - Université Laval, Quebec City, QC, Canada
34. Luxembourg Institute of Health, Luxembourg City, Luxembourg
35. Department of Neurological Surgery, Feinberg School of Medicine, Northwestern University, Chicago, Illinois.
36. Department of Cellular and Molecular Pathology, Leeds Teaching Hospital NHS Trust, St James’s University Hospital, Leeds, UK
37. Department of Neurology, Amsterdam University Medical Centers/VUmc, Amsterdam, The Netherlands
38. Department of Neurology, Columbia University Medical Center, New York, NY, USA
39. Department of Pathology and Cell Biology, Columbia University Medical Center, New York, NY, USA
40. Herbert Irving Comprehensive Cancer Center, Columbia University Medical Center, New York, NY, USA
41. Department of Radiology, Medical College of Wisconsin, Milwaukee WI, USA
42. Department of Oncologic Pathology, Dana-Farber Cancer Institute, Boston, MA, USA
43. Department of Pathology, Prince of Wales Hospital, Shatin, Hong Kong
44. Program for Mathematical Genomics, Columbia University Medical Center, New York, NY, USA
45. Department of Neurosurgery, The University of Texas MD Anderson Cancer Center, Houston, TX, USA
46. Institute of Neuropathology, Heinrich Heine University, Medical Faculty and University Hospital, Duesseldorf, Germany
47. Ivy Brain Tumor Center at the Barrow Neurological Institute, Phoenix, Arizona, USA
48. Department of Neurosurgery, Case Western Reserve University School of Medicine and University Hospitals of Cleveland, Cleveland, OH, USA
49. Erasmus MC, University Medical Centre Rotterdam, Rotterdam, The Netherlands
50. Division of Brain Tumor Translational Research, National Cancer Center Research Institute, Tokyo, Japan
51. Department of Neurology and Interdisciplinary Neuro-Oncology, University Hospital Tübingen and Hertie Institute for Clinical Brain Research, Tübingen, Germany
52. Department of Immunology, Duke University Medical Center, Durham, NC, USA
53. Department of Medical Oncology, Royal North Shore Hospital, St Leonards, Australia
54. Division of Neuropathology and Neurochemistry, Department of Neurology, Medical University of Vienna, Vienna, Austria
55. Department of Biostatistics and Bioinformatics, Duke University, Durham, NC, USA
56. Department of Mathematics, Duke University, Durham, NC, USA
57. Department of Neurosurgery, University of Toronto, Toronto, Ontario, Canada

## Acknowledgements

This work is supported by the National Institutes of Health under grant numbers, R01CA237208, R21NS114873 and P30CA034196 (R.G.W.V.), R01CA222146 (H.N., L.M.P., and I.D.), P30CA016672 and P50CA127001 (J.T.H., J.F.d.G., and K.D.A.), P30CA13148 (E.G.V.M.), U01CA242871, R01NS042645 and U24CA189523 (S.B.), R01CA218144 (P.S.L. and A.K.L.), P50CA190991 (J.X.), P50CA165962 and R01CA188228 (K.L.L.); the MD Anderson Moonshot (J.T.H., J.F.d.G., K.D.A., and D.R.O.); NCI-FCRDC contract 28XS100 (E.G.V.M.); the Leeds Hospitals Charity grant 9R11/14-11 (L.F.S.); the KWF Dutch Cancer Society project 11026 (P.W., M.C.M.K., M.S., and B.A.W.); the Department of Defense grant numbers CA170278 (H.N. and T.S.S.) and W81XWH1910246 (R.G.W.V); Roy and Diana Vagelos Precision Medicine Pilot Award (R.R. and J.Z.); and Strain for the Brain Milwaukee (P.S.L. and A.K.L.). This work was also supported by generous gifts from the Dabbiere family (R.G.W.V., J.F.C.). F.P.B. is supported by the JAX Scholar program and the National Cancer Institute (K99 CA226387). K.C.J. is the recipient of an American Cancer Society Fellowship (130984-PF-17-141-01-DMC). F.S.V. is supported by the JAX Scholar Program and a postdoctoral fellowship from The Jane Coffin Childs Memorial Fund for Medical Research.

## Author Contributions

Conceptualization, F.S.V., H.N., A.I., R.G.W.V.; Methodology, F.S.V.; Formal Analysis, F.S.V., K.C.J., N.A.; Investigation, F.S.V., K.C.J.; Resources, F.S.V., K.C.J., F.P.B., H.K., K.D.A., S.B., J.S.B.-S., C.B., A.B., J.M.C., J.F.C., I.D., J.F.d.G., H.K.G., A.C.H., C.H., J.T.H., A.I., P.S.L., K.L.L., A.K.L., T.M.M., A.M.M., M.P.N., H.K.N., S.P.N., D.R.O., S.H.P., J.J.P., L.M.P., G.Rao., G.Reifenberger., A.E.S., J.M.S., L.F.S., H.S., E.G.V.M., A.M.E.W., M.W., P.W., A.W. Data curation, F.S.V., K.C.J., T.E.W., T.M.M., T.S.S., F.P.B., H.K., N.A., I.D., S.B.A., A.C.H., M.K., M.C.M.K., L.M.P., D.R.O., L.F.S., C.W., M.W. Writing- Original draft, F.S.V., R.G.W.V.; Writing- Review & Editing, F.S.V., K.C.J., T.M.M., F.P.B., I.D., K.A., S.B., J.S.B.-S., H.G., P.V.G., M.K., E.K., J.T.H., S.C.S., G.T., E.G.V.M., A.M.E.W., C.W., T.W., M.W., P.W., A.W., J.X., L.F.S., L.M.P., H.N., A.I., R.G.W.V. Funding acquisition, R.G.W.V.; Supervision, H.N., A.I., R.G.W.V. All co-authors and contributors discussed the results and commented on the manuscript.

---

**Figure S1.**
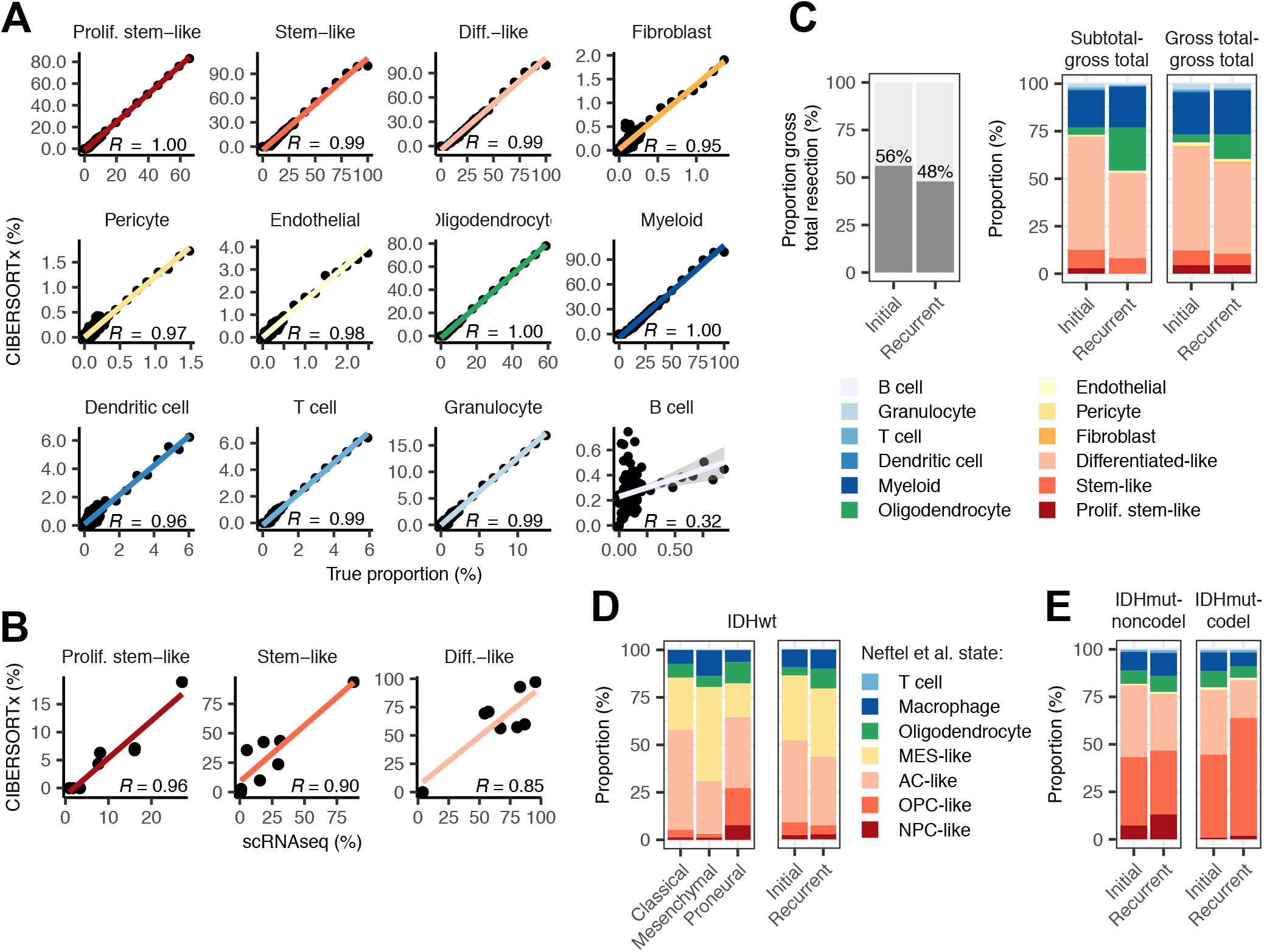
Validation of deconvolution results and IDH-wild-type-specific cell state profiles. Related to Figure 1. (A) Scatterplots depicting the association between the true proportion and the CIBERSORTx-inferred proportion for each cell state in gene expression profiles from synthetic mixtures composed of different combinations of single cells. (B) Scatterplots depicting the association between the proportion of each malignant cell state determined from single-cell RNAseq and the non-malignant cell-adjusted malignant cell state proportion inferred from CIBERSORTx applied to each sample’s respective bulk tumor RNAseq profile. In all plots, Pearson correlation coefficients are indicated. (C) Left: Stacked bar plot indicating the proportion of samples of IDH-wild-type tumors that underwent a gross total resection at each timepoint. Right: The average proportions of each cell state for tumors that underwent a subtotal resection at initial and a gross total resection at recurrence (Subtotal-gross total) and tumors that underwent a gross total resection at both time points (Gross total-gross total). (D) Left: The average Neftel et al. cell state composition of each bulk transcriptional subtype for all initial IDH-wild-type GLASS tumors. Right: The average Neftel et. al cell state composition of initial and recurrent IDH-wild-type tumors. (E) The average cell state composition of initial and recurrent IDH-mutant tumors stratified by 1p/19q co-deletion status. Colors in (E) are identical to those used in (C).

**Figure S2.**
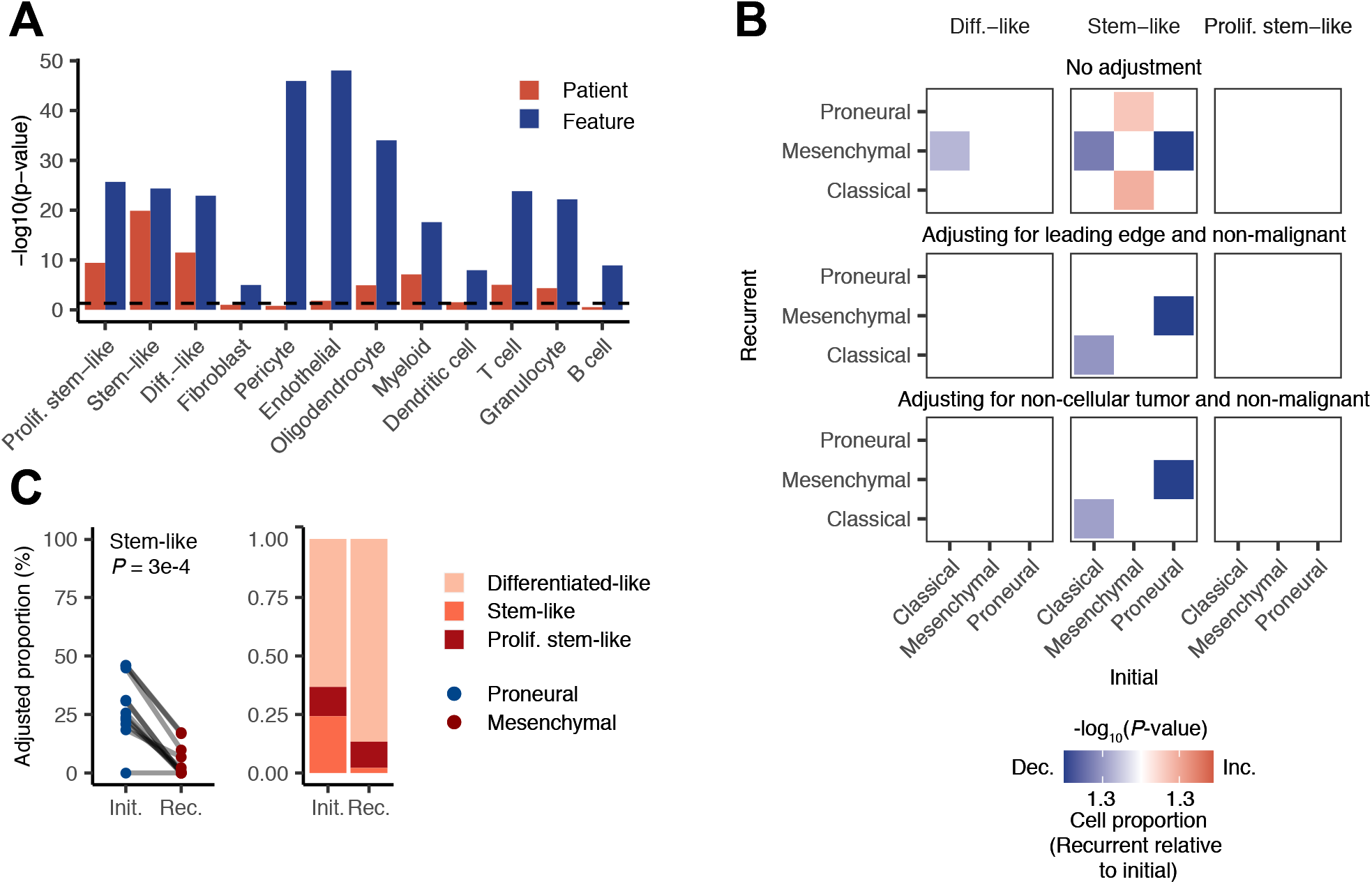
Relationship between bulk subtype switching and cell state changes after adjusting for histological feature composition. Related to Figure 2. (A) Bar plot depicting the -log_10_ *P*-value from a two-way ANOVA test measuring whether the fractions of each cell state in a sample associate with the patient the sample was derived from (red bar) and the feature the sample represents (blue bar). Dotted line corresponds to *P* = 0.05 (B) Heatmaps depicting the significance of the changes in each malignant cell state between initial and recurrent tumors undergoing the indicated subtype transition. The initial subtype is indicated in the columns and the recurrent subtype is indicated in the rows. Each row of heatmaps reflects a different histological feature adjustment. Colors represent the -log_10_(*P*-value) from a paired t-test, with increases at recurrence colored in red, decreases colored in blue, and *P*-values > 0.05 colored white. (C) Left: Ladder plot depicting the change in the adjusted stem-like cell proportion between paired initial and recurrent tumors undergoing a proneural-to-mesenchymal transition. Right: The average adjusted proportions for malignant cells for the tumor pairs outlined on the left. Malignant cell proportions were adjusted for the presence of non-malignant cells as well as all non-cellular tumor features.

**Figure S3.**
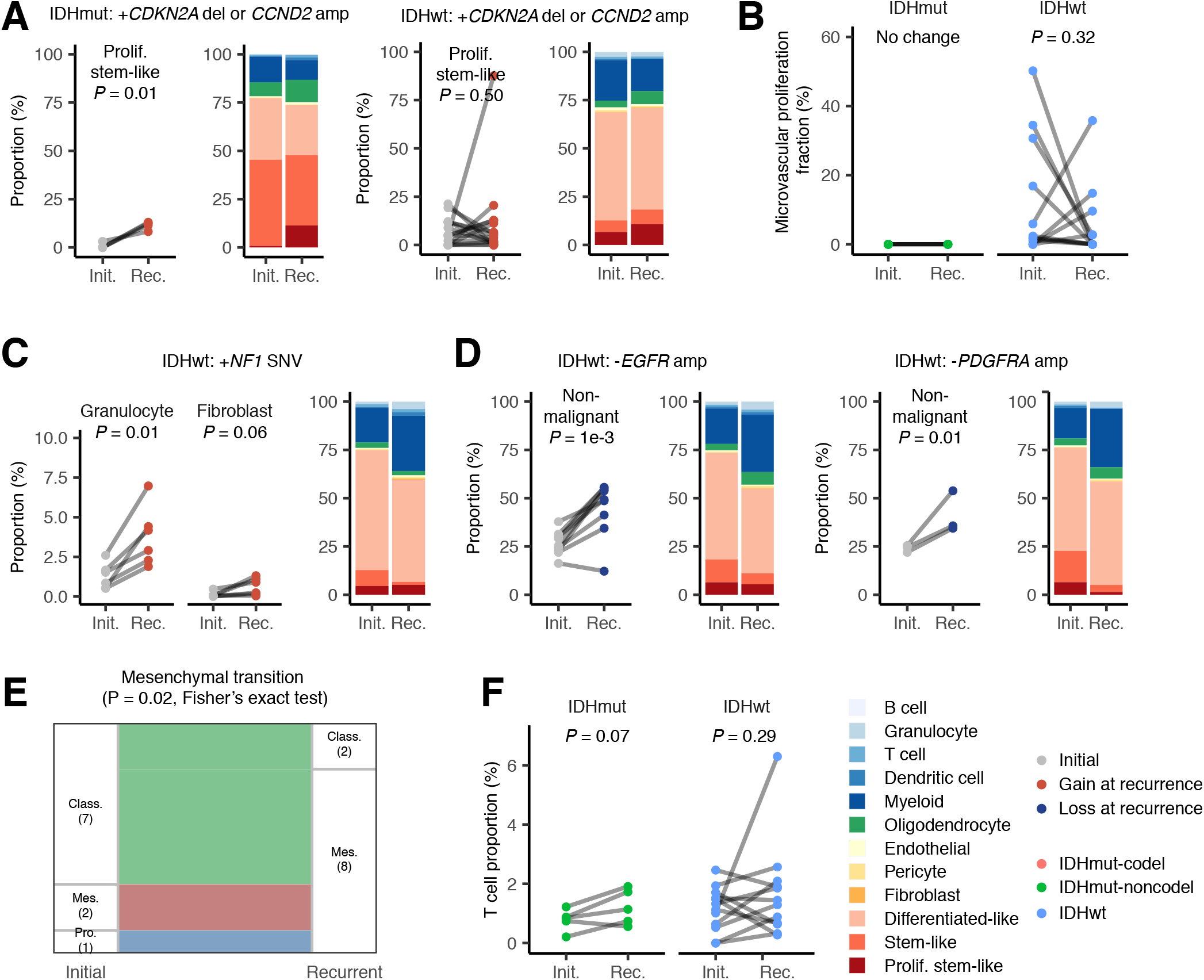
Cell state composition changes associated with the acquisition and loss of somatic alterations. Related to Figure 3. (A) Cell state differences in tumors that acquired *CDNK2A* deletions or *CCND2* amplifications. Panel is split into IDH-mutant and IDH-wild-type tumors. Ladder plots depict the change in the proliferating stem-like cell proportion between paired initial and recurrent tumors that acquired these alterations. Stacked bar plots depict the average proportions of each cell state for the tumor pairs in the ladder plots. (B) Ladder plots depicting the difference in microvascular proliferation fraction in IDH-mutant and IDH-wild-type tumors that underwent hypermutation at recurrence. (C) Left: Ladder plots depicting the change in granulocyte and fibroblast fractions in IDH-wild-type tumors that acquired mutations in *NF1* at recurrence. Right: The average proportions of each cell state for the tumor pairs in the ladder plots. (D) Non-malignant cell state differences in IDH-wild-type tumors that lost *EGFR* or *PDGFRA* amplifications at recurrence. Panel is split by alteration. Ladder plots depict the change in the non-malignant cell state proportion between paired initial and recurrent tumors while stacked bar plots depict the average proportions of each cell state for these tumors. (E) Sankey plot indicating whether the highest scoring transcriptional subtype changed at recurrence for the tumors depicted in (D). Each color reflects the transcriptional subtype in the initial tumors. Numbers in parentheses indicate number of samples. (F) Ladder plots depicting the difference in T cell fraction in IDH-mutant and IDH-wild-type tumors that underwent hypermutation at recurrence. In all figures, *P*- values were calculated using a paired t-test unless otherwise noted.

**Figure S4.**
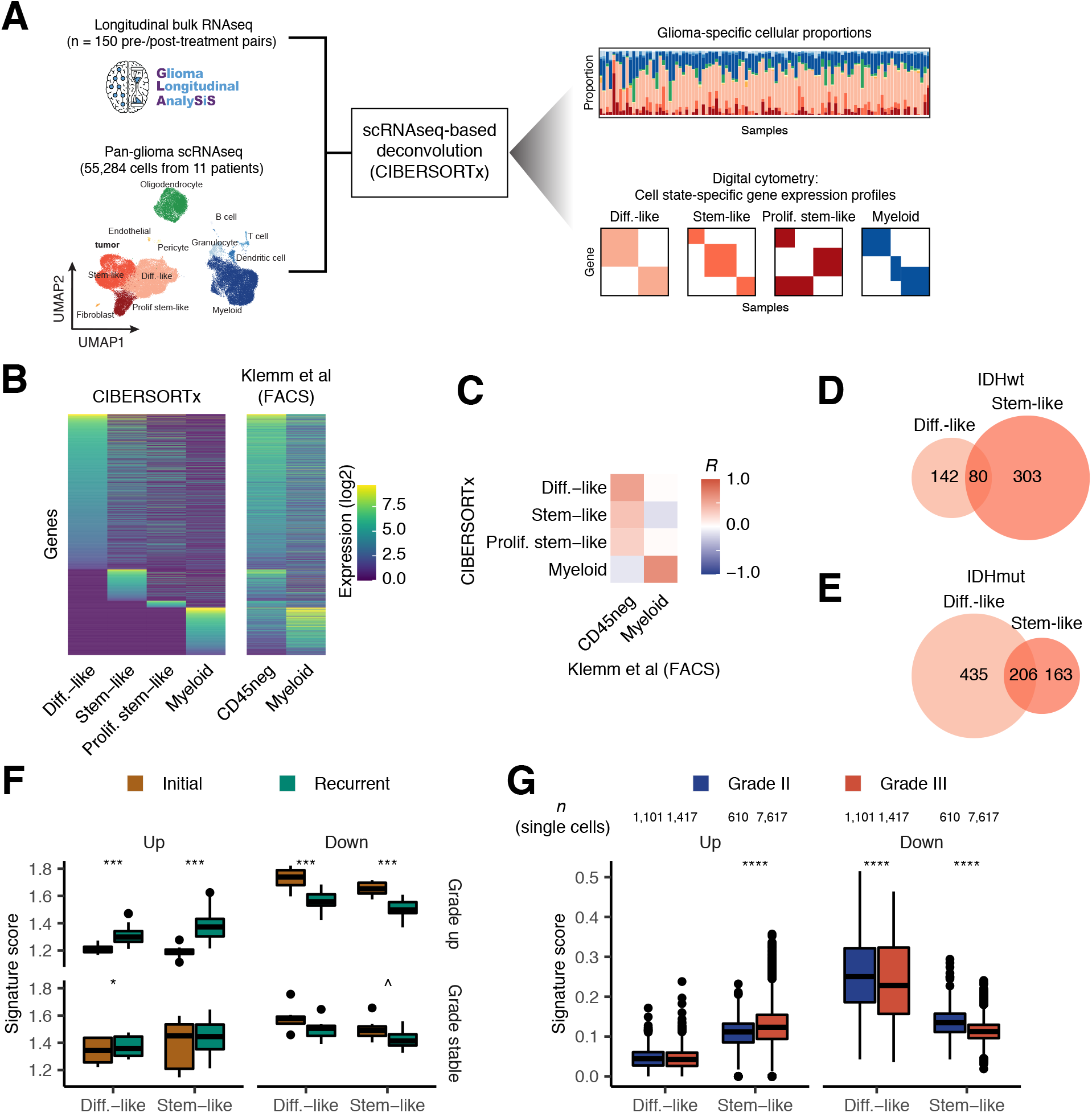
Validation and differential expression analysis of cell state-specific gene expression profiles. Related to Figure 4. (A) Schema for single-cell RNAseq-based deconvolution of cell state-specific gene expression profiles. (B) Left: Heatmap depicting the relationship between the CIBERSORTx-inferred gene expression profiles and gene expression profiles from analogous cell types from a FACS-purified ground truth dataset (Klemm et al.). In the CD45neg column in the Klemm et al. heatmap, which represents a composite gene expression profile from the non-immune cells purified from a collection of glioma tumors, gene expression patterns from all three malignant cell states can be observed. Right: Heatmap depicting the correlation coefficients between each CIBERSORTx-inferred cell state-specific gene expression profile and the gene expression profiles from the FACS-purified ground truth dataset. (D) Venn diagram depicting the overlap between the genes the differentiated-like and stem-like cell states differentially express in initial versus recurrent IDH-wild-type tumors. (E) Venn diagram depicting the overlap between the genes the differentiated-like and stem-like cell states differentially express in initial versus recurrent IDH-mutant tumors. (F) Boxplot depicting the average signature expression in the analogous cell state-specific gene expression profiles for each IDH-mutant tumor pair in GLASS. Comparisons are stratified based on whether the tumor pair was grade stable or exhibited a grade increase at recurrence. *** indicates Wilcoxon signed rank test *P*-value < 1e-3, * indicates *P* < 0.05, and ^ indicates *P* < 0.10. (G) Boxplot depicting the average signature expression in single cells of the indicated malignant cell states from grade II and grade III. **** indicates Wilcoxon rank-sum test *P*-value < 1e-5.

**Figure S5.**
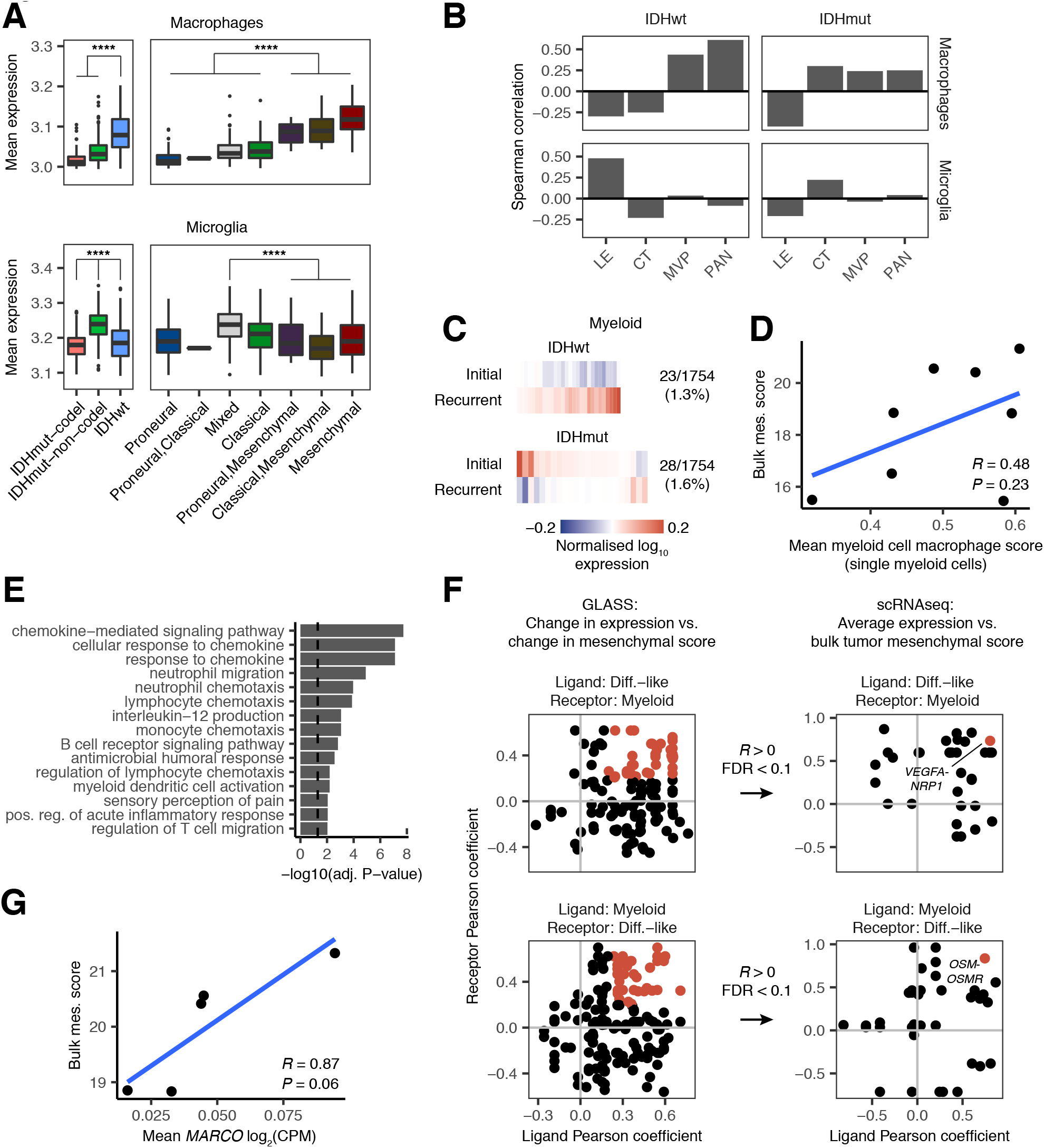
Characterization of the mesenchymal myeloid signature and identification of candidate ligand-receptor interactions in mesenchymal glioma. Related to Figure 5. (A) Boxplots depicting the average macrophage and microglia gene expression signatures in CIBERSORTx-inferred myeloid-specific gene expression profiles from TCGA. Samples are stratified by IDH and 1p/19q co-deletion status (left) and bulk transcriptional subtype (right). **** indicates Wilcoxon rank-sum test *P*-value < 1e-5. (B) Bar plots depicting the Spearman correlation coefficients measuring the association between the myeloid-specific expression scores for the macrophage and microglia signatures versus the presence of the four Ivy GAP histological features in TCGA. The features measured were leading edge (LE), cellular tumor (CT), microvascular proliferation (MVP), and pseudopalisading cells around necrosis (PAN). (C) Heatmaps depicting the average normalized log_10_ expression level of genes that were differentially expressed between myeloid cell states from initial and recurrent IDH-wild-type and IDH-mutant tumors in GLASS that did not undergo a subtype switch. Fractions on the right of each plot indicate the number of differentially expressed genes (numerator) out of the number of genes inferred for that cell state’s profile in GLASS using CIBERSORTx (denominator). (D) Scatterplot depicting the association between the mean blood-derived macrophage signature expression in single myeloid cells and the mesenchymal subtype score calculated from bulk RNAseq for each patient. (E) Bar plot depicting the -log_10_(adjusted *P*-value) from a GO enrichment analysis for the genes in the mesenchymal myeloid signature. (F) Analysis of ligand-receptor interactions between differentiated-like malignant cells and myeloid cells. Left plots depict the Pearson correlation coefficients from analyses comparing the change in expression of a ligand or receptor from the indicated cell state versus the change in bulk mesenchymal score over time in IDH-wild-type GLASS samples. All ligand-receptor pairs that exhibited an *R* > 0 and an FDR < 0.1 are highlighted in red and were included in the right plot. Right plots depict single-cell analyses measuring how the average expression of a ligand or receptor in single cells of the indicated cell state associates with the tumor’s bulk mesenchymal score in IDH-wild-type tumors. Red points indicate the ligand-receptor pair with the highest average correlation. (G) Scatterplot depicting the association between the mean expression of *MARCO2* in single myeloid cells and the mesenchymal subtype score calculated from bulk RNAseq for each patient.

**Figure S6.**
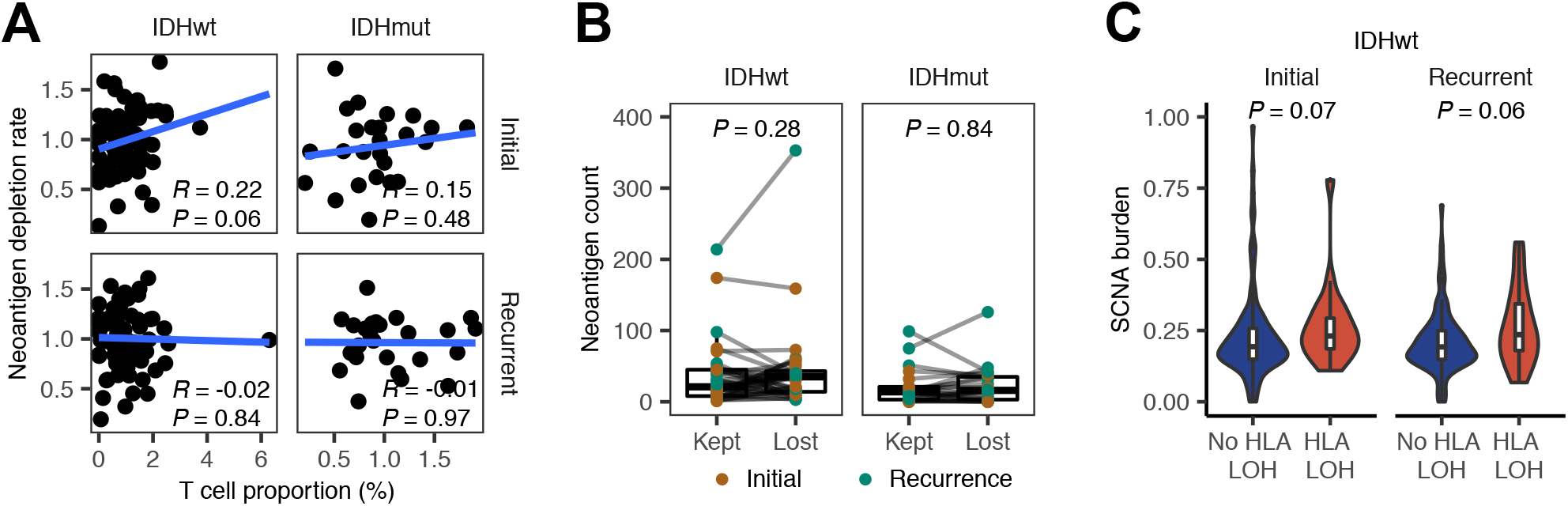
Analysis of neoantigen-mediated T cell selection in glioma. Related to Figure 6. (A) Scatterplots depicting the association between the T cell proportion and the neoantigen depletion rate in initial and recurrent GLASS samples. (B) Box and ladder plots depicting the difference in the number of neoantigens binding to the kept and lost allele. Points are colored based on whether the sample was an initial or recurrent tumor. *P*-values were calculated using the Wilcoxon signed-rank test. (C) Violin plots depicting the distribution of the somatic copynumber alteration burden in initial and recurrent IDH-wild-type GLASS samples that did and did not exhibit HLA LOH. *P*-values were calculated using the Wilcoxon rank-sum test.

## Methods

### GLASS Datasets

Datasets added to GLASS came from both published and unpublished sources (**Table S1**). Collectively, the newly added data consisted of exomes from 83 glioma samples (40 patients) and RNA-sequencing data from 351 samples (184 patients).

Newly generated whole exome data and RNAseq data was collected for a cohort of frozen samples from Henry Ford Health System. From each sample, DNA and RNA was simultaneously extracted using the AllPrep DNA/RNA Mini Kit from Qiagen (#80204). Exon capture was then performed using the Agilent’s SureSelect XT Low-Input Reagent Kit and the V6 + COSMIC capture library and the resulting reads were subjected to 150 base pair paired-end sequencing at the University of Southern California using an Illumina NovaSeq 6000. RNA from these tissues was processed and sequenced at Psomagen. New RNAseq data was also generated for cohorts coming from Case Western Reserve University, the Chinese University of Hong Kong, and MD Anderson Cancer Center. For Case Western Reserve University, RNA from frozen tissues was processed at Tempus (Chicago, IL) using the Tempus xO assay and then sequencing using an Illumina HiSeq 4000 platform. For the Chinese University of Hong Kong cohort, RNAseq libraries were prepared with the KAPA Stranded mRNAseq kit (Roche) per manufacturer’s instructions and then sequenced at The Jackson Laboratory for Genomic Medicine using an Illumina HiSeq4000 platform generating paired end reads of 75 base pairs. For the MD Anderson cohort, purified double-stranded cDNA generated from 150 ng of formalin-fixed paraffin-embedded (FFPE) sample-derived RNA was prepared using the NuGEN Ovation RNAseq System and subjected to paired-end sequencing using a HiSeq 2000 or HiSeq 2500 Sequencing System.

The remaining datasets were generated as described in their respective publications. For most of these cohorts, whole exome and/or whole genome sequencing data were downloaded and processed as described during creation of the initial GLASS dataset (Barthel et al., 2019). RNAseq fastq files from the Samsung Medical Center (SM) cohort were delivered via hard disk and are available to download from the European Genome-Phenome Archive (EGA) under accession numbers EGAS00001001041 and EGAS00001001880 (Kim et al., 2015b; Wang et al., 2016). RNAseq bam files for the original Henry Ford Health System (HF) and the University of California San Francisco (SF) cohorts were downloaded from EGA under accession numbers EGAS00001001033 and EGAS00001001255, respectively, and converted to fastq files for subsequent processing using bedtools (Kim et al., 2015a; Mazor et al., 2015). RNAseq fastq files for the University of Leeds (LU) cohort were downloaded from EGA under accession number EGAS00001003790 (Droop et al., 2018). For the first Columbia cohort (CU-R), which consisted of samples originally collected from the Istituto Neurologico C. Besta, RNAfastq files were delivered via hard disk and are available to download at the Sequencing Read Archive (SRA) under BioProject number PRJNA320312 (Wang et al., 2016). For the second Columbia cohort (CU-P), which featured samples that had been treated with immune checkpoint inhibitors, raw fastq reads for whole exome and RNAseq were obtained from SRA under BioProject number PRJNA482620 (Zhao et al., 2019). RNAseq fastq files from the Low Grade Glioma (LGG) and Glioblastoma Multiforme (GBM) projects in TCGA were obtained from the Genomic Data Commons legacy archive (https://portal.gdc.cancer.gov/legacy-archive/) (Brennan et al., 2013; Cancer Genome Atlas Research et al., 2015).

### Public Datasets

Processed RNAseq data from the TCGA glioma (GBMLGG) cohort was obtained from GDAC FireHose (RNAseqV2, RSEM). Normalized gene-level fragments per kilobase million (FPKM) for the Ivy Glioblastoma Atlas Project (Ivy GAP) dataset were obtained from the Ivy GAP website (https://glioblastoma.alleninstitute.org/static/download.html) (Puchalski et al., 2018). Processed single-cell data and associated metadata for a set of 28 IDH-wild-type glioblastomas processed using SmartSeq2 was obtained from the Broad Single Cell Portal (Study: Single cell RNA-seq of adult and pediatric glioblastoma; https://singlecell.broadinstitute.org/single_cell/study/SCP393/single-cell-rna-seq-of-adult-and-pediatric-glioblastoma) (Neftel et al., 2019). Raw count data and clinical annotation data from a set of glioma-derived cell populations purified using fluorescence activated cell sorting (FACS) was obtained from the Brain Tumor Immune Micro Environment (BrainTIME) portal and converted to counts per million (CPM) for downstream analysis (https://joycelab.shinyapps.io/braintime/) (Klemm et al., 2020).

### Whole exome and whole genome analysis

Whole exome and genome alignment, fingerprinting, variant detection, variant post-processing, mutation burden calculation, copy number segmentation, copy number calling, copy number-based purity, ploidy, HLA typing, and neoantigen calling were all performed using previously described pipelines that were developed during the initial GLASS data release (Barthel et al., 2019). Briefly, whole exome and whole genome reads were aligned to the b37 genome (human_g1k_v37_decoy) using BWA MEM 0.7.17 and pre-processed according to GATK Best Practices with GATK 4.0.10.1. Fingerprinting on the resulting files was performed using ‘CrosscheckFingerprints’ to confirm all readgroups from a given sample and all samples from a given patient match, with all mismatches being labelled and dropped from downstream analysis. Somatic mutations were called using GATK4.1 MuTect2. Hypermutation was defined for all recurrent tumors that had more than 10 mutations per megabase sequenced, as described previously (Barthel et al., 2019). Copy number segmentation and calling was performed according to GATK Best Practices as previously described. Copy number-based tumor purity and ploidy were determined using TITAN (Ha et al., 2014). Four-digit HLA class I types were determined from the normal bams for each sample using OptiType v1.3.2 (Szolek et al., 2014). Neoantigens were called from each patient’s somatic mutations and HLA types using pVACseq v4.0.10 (Hundal et al., 2016). Neoantigen depletion was calculated as described previously (Barthel et al., 2019). Loss of heterozygosity (LOH) for each sample’s HLA type was called from their respective matched tumor and normal bam files using LOHHLA run with default parameters and a coverage filter of 10 (https://bitbucket.org/mcgranahanlab/lohhla/) (McGranahan et al., 2017). HLA LOH was called if the estimated copy number for an allele using binning and B-allele frequency was < 0.5 and the *P*-value for allelic imbalance was < 0.05 (paired t-test).

### RNA preprocessing

To ensure each RNAseq file matched to the DNA and RNAseq files from their respective sample and patient, RNAseq fastq files were aligned to the b37 genome using STARv2.7.5 and the resulting bams were then preprocessed using the same pipelines described for DNA sequencing (Barthel et al., 2019). Fingerprinting was then performed on each bam at the readgroup and patient levels using ‘CrosscheckFingerprints.’ For each patient-level comparison, each RNA bam was compared to all other RNA and DNA bams coming from the same patient. All mismatches were labelled and dropped from downstream analysis.

RNAseq fastq files were pre-processed with fastp v0.20.0. Transcripts per million (TPM) values were then calculated from each sample’s set pre-processed files using kallisto v0.46.0 inputted with an index file built from the Ensemblv75 reference transcriptome. Strand-specific library preparation information was obtained for each sample from the source provider or using STARv2.7.5 quantMode set with the ‘GeneCounts’ parameter. The resulting TPM values for each sample were combined into a transcript expression matrix for downstream analysis. To create a gene expression matrix, transcript TPM values were collapsed and summed by their respective gene symbols.

### Quality control

For DNA samples to be included in longitudinal downstream analyses, two samples from a given patient had to pass a previously described quality control process based on fingerprinting, coverage, copy number variation, and clinical annotation criteria (Barthel et al., 2019). The resulting set of 243 whole exome or whole genome tumor pairs, known as the “gold set”, was used in all downstream DNA-only analyses. For RNA samples to be included in longitudinal downstream analyses, two samples from a given patient had to pass a patient-level fingerprinting filter that ensured that the RNA samples matched each other and the patient’s respective DNA samples if available, as well as a clinical annotation filter. The resulting set of 150 RNAseq pairs, known as the “RNA silver set”, was used in all downstream RNA-only analyses. Across the gold set and the RNA silver set, there were 101 tumor pairs that had DNA and RNA from the same sample at both timepoints. This overlapping set of pairs, known as the “platinum set”, was used in all downstream analyses that integrated DNA and RNA data.

### Bulk transcriptional subtype classification

Bulk transcriptional subtyping was performed on each GLASS or TCGA sample’s processed RNAseq profile using the “ssgsea.GBM.classification” R package (Wang et al., 2017). This method outputs an enrichment score quantifying the representation each of the three bulk glioma subtypes in a sample as well as a *P*-value indicating the significance of this representation. For each sample, the subtype with the lowest *P*-value was designated as that sample’s bulk transcriptional subtype. In cases where there were ties between subtypes, the subtype with the highest enrichment score was chosen.

### Joint single-cell and bulk RNAseq dataset

Single-cell and bulk RNA sequencing data were generated and processed as previously described (Johnson et al., 2020). Briefly, tumor surgical specimens were freshly collected, minced, and partitioned into single-cell and bulk fractions from the same tumor aliquot. The tissues aliquoted for single cell analyses were then mechanically and enzymatically dissociated using the Brain Tumor Dissociation Kit (P) according to the manufacturer’s protocol (Miltenyi Cat. No. 130-095-942). FACS was performed to select for viable single cells (Propidium Iodide-, Calcein+ singlets) and enrich for tumor cells by limiting the proportion of non-tumor cells (e.g., immune (CD45+) and endothelial (CD31+) cells). Sorted cells were then loaded on a 10X Chromium chip using the single-cell 3’ mRNA kit according to the manufacturer’s protocol (10X Genomics). A limitation of single-cell dissociation techniques is the exclusion of specific cell types, including neurons, that are found in glioma and surrounding tissue. Prior publications have estimated the neuronal content of central nervous system tumors to be less than 5% and therefore likely represent a minor non-malignant cell population in our dataset (Grabovska et al., 2020). The Cell Ranger pipeline (v3.0.2) was used to convert Illumina base call files to fastq files and align fastqs to hg19 10X reference genome (version 1.2.0) to be compatible with our bulk sequencing data. Data preprocessing and analysis was performed using the Scanpy package (1.3.7) (Wolf et al., 2018) with batch correction performed using BBKNN (Polanski et al., 2020). RNA was extracted for tissues with sufficient tissue and bulk RNAseq libraries were prepared with KAPA mRNA HyperPrep kit (Roche). Bulk RNA sequencing data was processed with the same pipeline as the GLASS samples.

### Deconvolution analyses

Cellular proportions and cell state-specific gene expression matrices were inferred from bulk RNAseq gene expression matrices using CIBERSORTx (Newman et al., 2019). Reference scRNAseq signature matrices were created from our internal 10x-derived scRNAseq dataset (Johnson et al., 2020) and a publicly available SmartSeq2-derived scRNAseq dataset (Neftel et al., 2019) using the ‘Create Signature Matrix’ module on the CIBERSORTx webserver (https://cibersortx.stanford.edu/) using default parameters and quantile normalization disabled. The Ivy GAP signature matrix was downloaded from a prior publication (Puchalski et al., 2018). The CIBERSORTx webserver currently recommends users input no more than 5,000 different single-cell profiles when creating their signature matrix (Steen et al., 2020). To meet this recommendation, our internal scRNAseq dataset, which is made up of 55,284 single cells, was randomly downsampled to 5,000 cells using the ‘sample’ command in R with the seed set to 11. The cells not included in signature matrix formation were then set aside for validation analyses.

Single-cell-derived cellular proportions and cell state-specific gene expression profiles were inferred from bulk RNAseq datasets using the CIBERSORTx High-Resolution docker container (https://hub.docker.com/r/cibersortx/hires) following CIBERSORTx instructions. For all runs, the bulk RNAseq dataset was input as the ‘mixture’ file and the respective signature matrix was input as the ‘sigmatrix’ file. For runs using our 10x-derived internal scRNAseq signatures, batch correction was done in ‘S-mode’ by setting the ‘rmbatchSmode’ parameter to TRUE, while for runs using SmartSeq2-derived scRNAseq signatures batch correction was done in ‘B-mode’ by setting the ‘rmbatchBmode’ parameter to TRUE. For each run, the inputted signature matrix’s respective CIBERSORTx-created “source gene expression profile” was input for batch correction. For all runs, the ‘subsetgenes’ parameter was set to a file containing the intersection of the gene symbols between the run’s respective source gene expression profile and the bulk RNAseq matrix that was being deconvoluted. For the run applying our internal scRNAseq dataset to the bulk GLASS RNAseq matrix, the ‘groundtruth’ parameter was set to a ground truth FACS-purified dataset that was generated as described below. Cellular proportions representing pre-created IvyGAP signatures were inferred using the ‘Impute Cell Fractions’ module on the CIBERSORTx webserver set to relative mode with quantile normalization and batch correction disabled and 100 permutations for significance analysis.

### Validation of cell state proportions and gene expression profiles

Cell state proportions derived from our internal scRNAseq dataset were validated using two approaches. In the first approach, synthetic mixtures were made using the single-cell gene expression profiles that had been left out of signature creation. Each synthetic mixture represented the average expression profile of 5,000 single cells where the number of cells of one cell state were manually set and the remaining cells were randomly sampled. Each cell state had its level manually set in 11 mixtures, where it represented 0% of the cells in the first mixture and then increased in 10% increments until reaching 100% in the final mixture. In cases where there were fewer than 5,000 single cells of a given cell state, making 100% representation not possible, the preset proportion instead represented the percent of available cells of that cell state rather than the percent of cells in the mixture. Each synthetic mixture had its true proportions recorded and the resulting mixtures were input into CIBERSORTx for deconvolution. Comparisons of the true and inferred proportions were then performed through correlation analysis. In the second approach, the cell state proportions inferred from bulk RNAseq data were compared to the cell state proportions quantified by scRNAseq for each sample in our internal scRNAseq dataset. Samples in this dataset were enriched for CD45^-^ cells via FACS and therefore precluded true cell state abundance when considering both malignant and non-malignant cells. To address this, comparisons were restricted to the relative proportions of each malignant cell state. Non- malignant cell proportions were removed, and malignant cells proportions were then renormalized so that the sum of each malignant cell state proportion in each sample added up to 1.

Concordance between CIBERSORTx-inferred cell state-specific gene expression profiles and a ground truth set of FACS-purified gene expression profiles was assessed using the ‘groundtruth’ parameter in CIBERSORTx. The ground truth dataset used in this step was generated from a previously released glioma dataset (Klemm et al., 2020) by collapsing all glioma-derived CD45^-^ profiles into an average CD45^-^ profile and all glioma-derived macrophage/microglia profiles into an average myeloid cell profile. This dataset was input into CIBERSORTx using the ‘groundtruth’ parameter during the run applying our internal scRNAseq signature matrix to the GLASS bulk RNAseq dataset. The resulting quality control files output during this run, primarily “SM_GEPs_HeatMap.txt”, were then used to perform correlation analyses assessing the similarity between the inferred malignant cell and myeloid profiles and the ground truth profiles.

### Analysis of cell state-specific gene expression profiles

To facilitate downstream analyses on each CIBERSORTx-inferred cell state-specific gene expression profile, each of the resulting expression matrices were log10-transformed and all genes that could not be imputed or had a variance of 0 across the dataset were removed. For each cell state-specific gene expression matrix, Wilcoxon signed-rank tests were used to determine the differentially expressed genes between initial and recurrent tumors and the resulting *P*-values were corrected for multiple testing using the Benjamini-Hochberg procedure. Signature scores in cell state-specific gene expression profiles and single-cell RNAseq profiles were defined as the average expression of the genes in the signature. In cases where the expression of some of the genes in the signature could not be determined, the intersection of the signature and the available genes was taken when calculating the signature score. For GO enrichment analyses on signatures derived from cell state-specific gene expression profiles, the background gene set only included the genes CIBERSORTx was able to impute for the cell state from which the signature was derived.

### Histological feature adjustment

For analyses examining how histological features influenced subtype switching, a tumor sample’s cell state composition profile was adjusted to remove cell states that could be attributed to a specific histological feature. To do this, the tumor sample’s proportion of a given histological feature was multiplied by the average proportion of each cell state from all samples of that feature in Ivy GAP. These numbers were then subtracted from their respective cell state’s proportion in the tumor sample and the resulting profile was then renormalized so that all proportions summed to 1. In cases where the new cell state proportion was less than 0, the value was set to 0 before renormalization.

### Statistical analysis

All data analyses were conducted in R 3.6.1 and PostgreSQL 10.6. GO enrichment analyses were performed using the “classic” algorithm in the R package “topGO” v2.38.1. When comparing variables between groups, t-tests were used for cell state proportions while non-parametric tests were used for all other variables (i.e., gene expression, signature score, neoantigen number). Clinical variables used throughout the study were defined as previously described in the Supplementary Information of the original GLASS study (Barthel et al., 2019).

### Code and data availability

All custom scripts, pipelines, and code used in figure creation will be made available at the time of publication on the project’s Github page. Processed data for the GLASS consortium is available on Synapse (https://www.synapse.org/#!Synapse:syn21589818) and will be publicly available on November 9, 2021.

## References

Bakas, S., Akbari, H., Pisapia, J., Martinez-Lage, M., Rozycki, M., Rathore, S., Dahmane, N., O’Rourke, D.M., and Davatzikos, C. (2017). In Vivo Detection of EGFRvIII in Glioblastoma via Perfusion Magnetic Resonance Imaging Signature Consistent with Deep Peritumoral Infiltration: The phi-Index. Clin Cancer Res 23, 4724–4734.

Bakas, S., Ormond, D.R., Alfaro-Munoz, K.D., Smits, M., Cooper, L.A.D., Verhaak, R., and Poisson, L.M. (2020). iGLASS: imaging integration into the Glioma Longitudinal Analysis Consortium. Neuro Oncol 22, 1545–1546.

Barthel, F.P., Johnson, K.C., Varn, F.S., Moskalik, A.D., Tanner, G., Kocakavuk, E., Anderson, K.J., Abiola, O., Aldape, K., Alfaro, K.D., et al. (2019). Longitudinal molecular trajectories of diffuse glioma in adults. Nature 576, 112–120.

Bhaduri, A., Di Lullo, E., Jung, D., Muller, S., Crouch, E.E., Espinosa, C.S., Ozawa, T., Alvarado, B., Spatazza, J., Cadwell, C.R., et al. (2020). Outer Radial Glia-like Cancer Stem Cells Contribute to Heterogeneity of Glioblastoma. Cell Stem Cell 26, 48–63 e46.

Bhat, K.P.L., Balasubramaniyan, V., Vaillant, B., Ezhilarasan, R., Hummelink, K., Hollingsworth, F., Wani, K., Heathcock, L., James, J.D., Goodman, L.D., et al. (2013). Mesenchymal differentiation mediated by NF-kappaB promotes radiation resistance in glioblastoma. Cancer Cell 24, 331–346.

Binder, Z.A., Thorne, A.H., Bakas, S., Wileyto, E.P., Bilello, M., Akbari, H., Rathore, S., Ha, S.M., Zhang, L., Ferguson, C.J., et al. (2018). Epidermal Growth Factor Receptor Extracellular Domain Mutations in Glioblastoma Present Opportunities for Clinical Imaging and Therapeutic Development. Cancer Cell 34, 163–177 e167.

Brennan, C.W., Verhaak, R.G., McKenna, A., Campos, B., Noushmehr, H., Salama, S.R., Zheng, S., Chakravarty, D., Sanborn, J.Z., Berman, S.H., et al. (2013). The somatic genomic landscape of glioblastoma. Cell 155, 462–477.

Cancer Genome Atlas Research, N., Brat, D.J., Verhaak, R.G., Aldape, K.D., Yung, W.K., Salama, S.R., Cooper, L.A., Rheinbay, E., Miller, C.R., Vitucci, M., et al. (2015). Comprehensive, Integrative Genomic Analysis of Diffuse Lower-Grade Gliomas. N Engl J Med 372, 2481–2498.

Castellan, M., Guarnieri, A., Fujimura, A., Zanconato, F., Battilana, G., Panciera, T., Sladitschek, H.L., Contessotto, P., Citron, A., Grilli, A., et al. (2021). Single-cell analyses reveal YAP/TAZ as regulators of stemness and cell plasticity in Glioblastoma. Nat Cancer 2, 174–188.

Ceccarelli, M., Barthel, F.P., Malta, T.M., Sabedot, T.S., Salama, S.R., Murray, B.A., Morozova, O., Newton, Y., Radenbaugh, A., Pagnotta, S.M., et al. (2016). Molecular Profiling Reveals Biologically Discrete Subsets and Pathways of Progression in Diffuse Glioma. Cell 164, 550–563.

Consortium, G. (2018). Glioma through the looking GLASS: molecular evolution of diffuse gliomas and the Glioma Longitudinal Analysis Consortium. Neuro Oncol 20, 873–884.

Couturier, C.P., Ayyadhury, S., Le, P.U., Nadaf, J., Monlong, J., Riva, G., Allache, R., Baig, S., Yan, X., Bourgey, M., et al. (2020). Single-cell RNA-seq reveals that glioblastoma recapitulates a normal neurodevelopmental hierarchy. Nat Commun 11, 3406.

Darmanis, S., Sloan, S.A., Croote, D., Mignardi, M., Chernikova, S., Samghababi, P., Zhang, Y., Neff, N., Kowarsky, M., Caneda, C., et al. (2017). Single-Cell RNA-Seq Analysis of Infiltrating Neoplastic Cells at the Migrating Front of Human Glioblastoma. Cell Rep 21, 1399–1410.

Droop, A., Bruns, A., Tanner, G., Rippaus, N., Morton, R., Harrison, S., King, H., Ashton, K., Syed, K., Jenkinson, M.D., et al. (2018). How to analyse the spatiotemporal tumour samples needed to investigate cancer evolution: A case study using paired primary and recurrent glioblastoma. Int J Cancer 142, 1620–1626.

Eckel-Passow, J.E., Lachance, D.H., Molinaro, A.M., Walsh, K.M., Decker, P.A., Sicotte, H., Pekmezci, M., Rice, T., Kosel, M.L., Smirnov, I.V., et al. (2015). Glioma Groups Based on 1p/19q, IDH, and TERT Promoter Mutations in Tumors. N Engl J Med 372, 2499–2508.

Fathi Kazerooni, A., Bakas, S., Saligheh Rad, H., and Davatzikos, C. (2020). Imaging signatures of glioblastoma molecular characteristics: A radiogenomics review. J Magn Reson Imaging 52, 54–69.

Gangoso, E., Southgate, B., Bradley, L., Rus, S., Galvez-Cancino, F., McGivern, N., Guc, E., Kapourani, C.A., Byron, A., Ferguson, K.M., et al. (2021). Glioblastomas acquire myeloid-affiliated transcriptional programs via epigenetic immunoediting to elicit immune evasion. Cell.

Garofano, L., Migliozzi, S., Oh, Y.T., D’Angelo, F., Najac, R.D., Ko, A., Frangaj, B., Caruso, F.P., Yu, K., Yuan, J., et al. (2021). Pathway-based classification of glioblastoma uncovers a mitochondrial subtype with therapeutic vulnerabilities. Nat Cancer 2, 141–156.

Gill, B.J., Pisapia, D.J., Malone, H.R., Goldstein, H., Lei, L., Sonabend, A., Yun, J., Samanamud, J., Sims, J.S., Banu, M., et al. (2014). MRI-localized biopsies reveal subtype-specific differences in molecular and cellular composition at the margins of glioblastoma. Proc Natl Acad Sci U S A 111, 12550–12555.

Grabovska, Y., Mackay, A., O’Hare, P., Crosier, S., Finetti, M., Schwalbe, E.C., Pickles, J.C., Fairchild, A.R., Avery, A., Cockle, J., et al. (2020). Pediatric pan-central nervous system tumor analysis of immune-cell infiltration identifies correlates of antitumor immunity. Nat Commun 11, 4324.

Grasso, C.S., Giannakis, M., Wells, D.K., Hamada, T., Mu, X.J., Quist, M., Nowak, J.A., Nishihara, R., Qian, Z.R., Inamura, K., et al. (2018). Genetic Mechanisms of Immune Evasion in Colorectal Cancer. Cancer Discov 8, 730–749.

Ha, G., Roth, A., Khattra, J., Ho, J., Yap, D., Prentice, L.M., Melnyk, N., McPherson, A., Bashashati, A., Laks, E., et al. (2014). TITAN: inference of copy number architectures in clonal cell populations from tumor whole-genome sequence data. Genome Res 24, 1881–1893.

Hambardzumyan, D., and Bergers, G. (2015). Glioblastoma: Defining Tumor Niches. Trends Cancer 1, 252–265.

Herzog, B., Pellet-Many, C., Britton, G., Hartzoulakis, B., and Zachary, I.C. (2011). VEGF binding to NRP1 is essential for VEGF stimulation of endothelial cell migration, complex formation between NRP1 and VEGFR2, and signaling via FAK Tyr407 phosphorylation. Mol Biol Cell 22, 2766–2776.

Hundal, J., Carreno, B.M., Petti, A.A., Linette, G.P., Griffith, O.L., Mardis, E.R., and Griffith, M. (2016). pVAC-Seq: A genome-guided in silico approach to identifying tumor neoantigens. Genome Med 8, 11.

Jin, X., Kim, L.J.Y., Wu, Q., Wallace, L.C., Prager, B.C., Sanvoranart, T., Gimple, R.C., Wang, X., Mack, S.C., Miller, T.E., et al. (2017). Targeting glioma stem cells through combined BMI1 and EZH2 inhibition. Nat Med 23, 1352–1361.

Johnson, K.C., Anderson, K.J., Courtois, E.T., Barthel, F.P., Varn, F.S., Luo, D., Seignon, M., Yi, E., Kim, H., Estecio, M.R., et al. (2020). Single-cell multimodal glioma analyses reveal epigenetic regulators of cellular plasticity and environmental stress response. bioRxiv, 2020.2007.2022.215335. Nat Gen. In press.

Junk, D.J., Bryson, B.L., Smigiel, J.M., Parameswaran, N., Bartel, C.A., and Jackson, M.W. (2017). Oncostatin M promotes cancer cell plasticity through cooperative STAT3-SMAD3 signaling. Oncogene 36, 4001–4013.

Kim, H., Zheng, S., Amini, S.S., Virk, S.M., Mikkelsen, T., Brat, D.J., Grimsby, J., Sougnez, C., Muller, F., Hu, J., et al. (2015a). Whole-genome and multisector exome sequencing of primary and post-treatment glioblastoma reveals patterns of tumor evolution. Genome Res 25, 316–327.

Kim, J., Lee, I.H., Cho, H.J., Park, C.K., Jung, Y.S., Kim, Y., Nam, S.H., Kim, B.S., Johnson, M.D., Kong, D.S., et al. (2015b). Spatiotemporal Evolution of the Primary Glioblastoma Genome. Cancer Cell 28, 318–328.

Kim, Y., Varn, F.S., Park, S.H., Yoon, B.W., Park, H.R., Lee, C., Verhaak, R.G.W., and Paek, S.H. (2021). Perspective of mesenchymal transformation in glioblastoma. Acta Neuropathol Commun 9, 50.

Klemm, F., Maas, R.R., Bowman, R.L., Kornete, M., Soukup, K., Nassiri, S., Brouland, J.P., Iacobuzio-Donahue, C.A., Brennan, C., Tabar, V., et al. (2020). Interrogation of the Microenvironmental Landscape in Brain Tumors Reveals Disease-Specific Alterations of Immune Cells. Cell 181, 1643–1660 e1617.

Korber, V., Yang, J., Barah, P., Wu, Y., Stichel, D., Gu, Z., Fletcher, M.N.C., Jones, D., Hentschel, B., Lamszus, K., et al. (2019). Evolutionary Trajectories of IDH(WT) Glioblastomas Reveal a Common Path of Early Tumorigenesis Instigated Years ahead of Initial Diagnosis. Cancer Cell 35, 692–704 e612.

Kristensen, B.W., Priesterbach-Ackley, L.P., Petersen, J.K., and Wesseling, P. (2019). Molecular pathology of tumors of the central nervous system. Ann Oncol 30, 1265–1278.

Louis, D.N., Perry, A., Reifenberger, G., von Deimling, A., Figarella-Branger, D., Cavenee, W.K., Ohgaki, H., Wiestler, O.D., Kleihues, P., and Ellison, D.W. (2016). The 2016 World Health Organization Classification of Tumors of the Central Nervous System: a summary. Acta Neuropathol 131, 803–820.

Mang, A., Bakas, S., Subramanian, S., Davatzikos, C., and Biros, G. (2020). Integrated Biophysical Modeling and Image Analysis: Application to Neuro-Oncology. Annu Rev Biomed Eng 22, 309–341.

Mao, P., Joshi, K., Li, J., Kim, S.H., Li, P., Santana-Santos, L., Luthra, S., Chandran, U.R., Benos, P.V., Smith, L., et al. (2013). Mesenchymal glioma stem cells are maintained by activated glycolytic metabolism involving aldehyde dehydrogenase 1A3. Proc Natl Acad Sci U S A 110, 8644–8649.

Mazor, T., Pankov, A., Johnson, B.E., Hong, C., Hamilton, E.G., Bell, R.J.A., Smirnov, I.V., Reis, G.F., Phillips, J.J., Barnes, M.J., et al. (2015). DNA Methylation and Somatic Mutations Converge on the Cell Cycle and Define Similar Evolutionary Histories in Brain Tumors. Cancer Cell 28, 307–317.

McGranahan, N., Rosenthal, R., Hiley, C.T., Rowan, A.J., Watkins, T.B.K., Wilson, G.A., Birkbak, N.J., Veeriah, S., Van Loo, P., Herrero, J., et al. (2017). Allele-Specific HLA Loss and Immune Escape in Lung Cancer Evolution. Cell 171, 1259–1271 e1211.

Muller, S., Kohanbash, G., Liu, S.J., Alvarado, B., Carrera, D., Bhaduri, A., Watchmaker, P.B., Yagnik, G., Di Lullo, E., Malatesta, M., et al. (2017). Single-cell profiling of human gliomas reveals macrophage ontogeny as a basis for regional differences in macrophage activation in the tumor microenvironment. Genome Biol 18, 234.

Neftel, C., Laffy, J., Filbin, M.G., Hara, T., Shore, M.E., Rahme, G.J., Richman, A.R., Silverbush, D., Shaw, M.L., Hebert, C.M., et al. (2019). An Integrative Model of Cellular States, Plasticity, and Genetics for Glioblastoma. Cell 178, 835–849 e821.

Newman, A.M., Steen, C.B., Liu, C.L., Gentles, A.J., Chaudhuri, A.A., Scherer, F., Khodadoust, M.S., Esfahani, M.S., Luca, B.A., Steiner, D., et al. (2019). Determining cell type abundance and expression from bulk tissues with digital cytometry. Nat Biotechnol 37, 773–782.

Ochocka, N., Segit, P., Walentynowicz, K.A., Wojnicki, K., Cyranowski, S., Swatler, J., Mieczkowski, J., and Kaminska, B. (2021). Single-cell RNA sequencing reveals functional heterogeneity of glioma-associated brain macrophages. Nat Commun 12, 1151.

Osuka, S., Zhu, D., Zhang, Z., Li, C., Stackhouse, C.T., Sampetrean, O., Olson, J.J., Gillespie, G.Y., Saya, H., Willey, C.D., et al. (2021). N-cadherin upregulation mediates adaptive radioresistance in glioblastoma. J Clin Invest 131.

Patel, A.P., Tirosh, I., Trombetta, J.J., Shalek, A.K., Gillespie, S.M., Wakimoto, H., Cahill, D.P., Nahed, B.V., Curry, W.T., Martuza, R.L., et al. (2014). Single-cell RNA-seq highlights intratumoral heterogeneity in primary glioblastoma. Science 344, 1396–1401.

Phillips, H.S., Kharbanda, S., Chen, R., Forrest, W.F., Soriano, R.H., Wu, T.D., Misra, A., Nigro, J.M., Colman, H., Soroceanu, L., et al. (2006). Molecular subclasses of high-grade glioma predict prognosis, delineate a pattern of disease progression, and resemble stages in neurogenesis. Cancer Cell 9, 157–173.

Polanski, K., Young, M.D., Miao, Z., Meyer, K.B., Teichmann, S.A., and Park, J.E. (2020). BBKNN: fast batch alignment of single cell transcriptomes. Bioinformatics 36, 964–965.

Pombo Antunes, A.R., Scheyltjens, I., Lodi, F., Messiaen, J., Antoranz, A., Duerinck, J., Kancheva, D., Martens, L., De Vlaminck, K., Van Hove, H., et al. (2021). Single-cell profiling of myeloid cells in glioblastoma across species and disease stage reveals macrophage competition and specialization. Nat Neurosci 24, 595–610.

Puchalski, R.B., Shah, N., Miller, J., Dalley, R., Nomura, S.R., Yoon, J.G., Smith, K.A., Lankerovich, M., Bertagnolli, D., Bickley, K., et al. (2018). An anatomic transcriptional atlas of human glioblastoma. Science 360, 660–663.

Ramilowski, J.A., Goldberg, T., Harshbarger, J., Kloppmann, E., Lizio, M., Satagopam, V.P., Itoh, M., Kawaji, H., Carninci, P., Rost, B., et al. (2015). A draft network of ligand-receptor-mediated multicellular signalling in human. Nat Commun 6, 7866.

Richards, L.M., Whitley, O.K.N., MacLeod, G., Cavalli, F.M.G., Coutinho, F.J., Jaramillo, J.E., Svergun, N., Riverin, M., Croucher, D.C., Kushida, M., et al. (2021). Gradient of Developmental and Injury Response transcriptional states defines functional vulnerabilities underpinning glioblastoma heterogeneity. Nature Cancer 2, 157–173.

Rooney, M.S., Shukla, S.A., Wu, C.J., Getz, G., and Hacohen, N. (2015). Molecular and genetic properties of tumors associated with local immune cytolytic activity. Cell 160, 48–61.

Rosenthal, R., Cadieux, E.L., Salgado, R., Bakir, M.A., Moore, D.A., Hiley, C.T., Lund, T., Tanic, M., Reading, J.L., Joshi, K., et al. (2019). Neoantigen-directed immune escape in lung cancer evolution. Nature 567, 479–485.

Sa, J.K., Chang, N., Lee, H.W., Cho, H.J., Ceccarelli, M., Cerulo, L., Yin, J., Kim, S.S., Caruso, F.P., Lee, M., et al. (2020). Transcriptional regulatory networks of tumor-associated macrophages that drive malignancy in mesenchymal glioblastoma. Genome Biol 21, 216.

Schmitt, M.J., Company, C., Dramaretska, Y., Barozzi, I., Gohrig, A., Kertalli, S., Grossmann, M., Naumann, H., Sanchez-Bailon, M.P., Hulsman, D., et al. (2021). Phenotypic Mapping of Pathologic Cross-Talk between Glioblastoma and Innate Immune Cells by Synthetic Genetic Tracing. Cancer Discov 11, 754–777.

Steen, C.B., Liu, C.L., Alizadeh, A.A., and Newman, A.M. (2020). Profiling Cell Type Abundance and Expression in Bulk Tissues with CIBERSORTx. In Stem Cell Transcriptional Networks: Methods and Protocols, B.L. Kidder, ed. (New York, NY: Springer US), pp. 135-157.

Stupp, R., Mason, W.P., van den Bent, M.J., Weller, M., Fisher, B., Taphoorn, M.J., Belanger, K., Brandes, A.A., Marosi, C., Bogdahn, U., et al. (2005). Radiotherapy plus concomitant and adjuvant temozolomide for glioblastoma. N Engl J Med 352, 987–996.

Szolek, A., Schubert, B., Mohr, C., Sturm, M., Feldhahn, M., and Kohlbacher, O. (2014). OptiType: precision HLA typing from next-generation sequencing data. Bioinformatics 30, 3310–3316.

Tirosh, I., Venteicher, A.S., Hebert, C., Escalante, L.E., Patel, A.P., Yizhak, K., Fisher, J.M., Rodman, C., Mount, C., Filbin, M.G., et al. (2016). Single-cell RNA-seq supports a developmental hierarchy in human oligodendroglioma. Nature 539, 309–313.

Touat, M., Li, Y.Y., Boynton, A.N., Spurr, L.F., Iorgulescu, J.B., Bohrson, C.L., Cortes-Ciriano, I., Birzu, C., Geduldig, J.E., Pelton, K., et al. (2020). Mechanisms and therapeutic implications of hypermutation in gliomas. Nature 580, 517–523.

Venkataramani, V., Tanev, D.I., Strahle, C., Studier-Fischer, A., Fankhauser, L., Kessler, T., Korber, C., Kardorff, M., Ratliff, M., Xie, R., et al. (2019). Glutamatergic synaptic input to glioma cells drives brain tumour progression. Nature 573, 532–538.

Venkatesh, H.S., Johung, T.B., Caretti, V., Noll, A., Tang, Y., Nagaraja, S., Gibson, E.M., Mount, C.W., Polepalli, J., Mitra, S.S., et al. (2015). Neuronal Activity Promotes Glioma Growth through Neuroligin-3 Secretion. Cell 161, 803–816.

Venkatesh, H.S., Morishita, W., Geraghty, A.C., Silverbush, D., Gillespie, S.M., Arzt, M., Tam, L.T., Espenel, C., Ponnuswami, A., Ni, L., et al. (2019). Electrical and synaptic integration of glioma into neural circuits. Nature 573, 539–545.

Venkatesh, H.S., Tam, L.T., Woo, P.J., Lennon, J., Nagaraja, S., Gillespie, S.M., Ni, J., Duveau, D.Y., Morris, P.J., Zhao, J.J., et al. (2017). Targeting neuronal activity-regulated neuroligin-3 dependency in high-grade glioma. Nature 549, 533–537.

Venteicher, A.S., Tirosh, I., Hebert, C., Yizhak, K., Neftel, C., Filbin, M.G., Hovestadt, V., Escalante, L.E., Shaw, M.L., Rodman, C., et al. (2017). Decoupling genetics, lineages, and microenvironment in IDH-mutant gliomas by single-cell RNA-seq. Science 355.

Verhaak, R.G., Hoadley, K.A., Purdom, E., Wang, V., Qi, Y., Wilkerson, M.D., Miller, C.R., Ding, L., Golub, T., Mesirov, J.P., et al. (2010). Integrated genomic analysis identifies clinically relevant subtypes of glioblastoma characterized by abnormalities in PDGFRA, IDH1, EGFR, and NF1. Cancer Cell 17, 98–110.

Wang, J., Cazzato, E., Ladewig, E., Frattini, V., Rosenbloom, D.I., Zairis, S., Abate, F., Liu, Z., Elliott, O., Shin, Y.J., et al. (2016). Clonal evolution of glioblastoma under therapy. Nat Genet 48, 768–776.

Wang, L., Babikir, H., Muller, S., Yagnik, G., Shamardani, K., Catalan, F., Kohanbash, G., Alvarado, B., Di Lullo, E., Kriegstein, A., et al. (2019). The Phenotypes of Proliferating Glioblastoma Cells Reside on a Single Axis of Variation. Cancer Discov 9, 1708–1719.

Wang, Q., Hu, B., Hu, X., Kim, H., Squatrito, M., Scarpace, L., deCarvalho, A.C., Lyu, S., Li, P., Li, Y., et al. (2017). Tumor Evolution of Glioma-Intrinsic Gene Expression Subtypes Associates with Immunological Changes in the Microenvironment. Cancer Cell 32, 42–56 e46.

Weller, M., Weber, R.G., Willscher, E., Riehmer, V., Hentschel, B., Kreuz, M., Felsberg, J., Beyer, U., Loffler-Wirth, H., Kaulich, K., et al. (2015). Molecular classification of diffuse cerebral WHO grade II/III gliomas using genome- and transcriptome-wide profiling improves stratification of prognostically distinct patient groups. Acta Neuropathol 129, 679–693.

Wen, P.Y., Weller, M., Lee, E.Q., Alexander, B.M., Barnholtz-Sloan, J.S., Barthel, F.P., Batchelor, T.T., Bindra, R.S., Chang, S.M., Chiocca, E.A., et al. (2020). Glioblastoma in adults: a Society for Neuro-Oncology (SNO) and European Society of Neuro-Oncology (EANO) consensus review on current management and future directions. Neuro Oncol 22, 1073–1113.

Wolf, F.A., Angerer, P., and Theis, F.J. (2018). SCANPY: large-scale single-cell gene expression data analysis. Genome Biol 19, 15.

Xue, J., Schmidt, S.V., Sander, J., Draffehn, A., Krebs, W., Quester, I., De Nardo, D., Gohel, T.D., Emde, M., Schmidleithner, L., et al. (2014). Transcriptome-based network analysis reveals a spectrum model of human macrophage activation. Immunity 40, 274–288.

Yan, H., Parsons, D.W., Jin, G., McLendon, R., Rasheed, B.A., Yuan, W., Kos, I., Batinic-Haberle, I., Jones, S., Riggins, G.J., et al. (2009). IDH1 and IDH2 mutations in gliomas. N Engl J Med 360, 765–773.

Yee, P.P., Wei, Y., Kim, S.Y., Lu, T., Chih, S.Y., Lawson, C., Tang, M., Liu, Z., Anderson, B., Thamburaj, K., et al. (2020). Neutrophil-induced ferroptosis promotes tumor necrosis in glioblastoma progression. Nat Commun 11, 5424.

Yu, Y., Villanueva-Meyer, J., Grimmer, M.R., Hilz, S., Solomon, D.A., Choi, S., Wahl, M., Mazor, T., Hong, C., Shai, A., et al. (2021). Temozolomide-induced hypermutation is associated with distant recurrence and reduced survival after high-grade transformation of low-grade IDH-mutant gliomas. Neuro Oncol.

Yuan, J., Levitin, H.M., Frattini, V., Bush, E.C., Boyett, D.M., Samanamud, J., Ceccarelli, M., Dovas, A., Zanazzi, G., Canoll, P., et al. (2018). Single-cell transcriptome analysis of lineage diversity in high-grade glioma. Genome Med 10, 57.

Zhang, A.W., McPherson, A., Milne, K., Kroeger, D.R., Hamilton, P.T., Miranda, A., Funnell, T., Little, N., de Souza, C.P.E., Laan, S., et al. (2018). Interfaces of Malignant and Immunologic Clonal Dynamics in Ovarian Cancer. Cell 173, 1755–1769 e1722.

Zhao, J., Chen, A.X., Gartrell, R.D., Silverman, A.M., Aparicio, L., Chu, T., Bordbar, D., Shan, D., Samanamud, J., Mahajan, A., et al. (2019). Immune and genomic correlates of response to anti- PD-1 immunotherapy in glioblastoma. Nat Med 25, 462–469.

Zwanenburg, A., Vallieres, M., Abdalah, M.A., Aerts, H., Andrearczyk, V., Apte, A., Ashrafinia, S., Bakas, S., Beukinga, R.J., Boellaard, R., et al. (2020). The Image Biomarker Standardization Initiative: Standardized Quantitative Radiomics for High-Throughput Image-based Phenotyping. Radiology 295, 328–338.

